# Balancing stability, dynamics and kinetics in phase separation of intrinsically disordered proteins

**DOI:** 10.1101/2024.01.05.574441

**Authors:** Guoqing Zhang, Xiakun Chu

**Affiliations:** Advanced Materials Thrust, Function Hub; Green e Materials Laboratory; Guangzhou Municipal Key Laboratory of Materials Informatics, The Hong Kong University of Science and Technology (Guangzhou), Guangzhou, Guangdong 511400, China; Division of Life Science, The Hong Kong University of Science and Technology, Clear Water Bay, Hong Kong SAR 999077, China

## Abstract

Liquid-liquid phase separation is a ubiquitous molecular phenomenon that plays crucial roles in a multitude of essential cellular activities. Intrinsically disordered proteins (IDPs), which lack well-defined three-dimensional structures, are prevalent participants in phase separation due to their inherent potential for promoting multivalent binding–the major driving force for this process. Understanding the underlying mechanisms of phase separation is challenging, as phase separation is a complex process, involving numerous molecules and various types of interactions. Here, we used a simplified coarse-grained model of IDPs to investigate the thermodynamic stability of the dense phase, conformational properties of IDPs, chain dynamics and kinetic rates of forming condensates. We focused on the IDP system, in which the oppositely charged IDPs are maximally segregated, inherently possessing a high propensity for phase separation. By varying interaction strengths, salt concentrations and temperatures, we observed that IDPs in the dense phase exhibited highly conserved conformational characteristics, which are more extended than those in the dilute phase. This implies that condensate formation acts as a protective shield, enabling IDPs to maintain conformational ensemble with high resistance to the changes in interactions and environmental conditions. Although the chain motions and global conformational dynamics of IDPs in the condensates are slow due to the high viscosity, local chain flexibility at the short timescales is largely preserved with respect to that at the free state. Strikingly, we observed a non-monotonic relationship between interaction strengths and kinetic rates for forming condensates. As strong interactions of IDPs result in high stable condensates, our results suggest that the thermodynamics and kinetics of phase separation are decoupled and optimized by the speed-stability balance through underlying molecular interactions. Our findings contribute to the molecular-level understanding of phase separation and offer valuable insights into the developments of engineering strategies for precise regulation of biomolecular condensates.

## 1 Introduction

Increasing evidence revealed that many intrinsically disordered proteins (IDPs) can undergo liquid-liquid phase separation, where the protein-rich phase and the dilute phase coexist in solution [1, 2, 3, 4]. Phase separation is recognized to play crucial roles in various essential biological processes by forming membrane-less organelles [5, 6, 7, 8]. Despite being involved in diverse fundamental cellular processes, these high-order phase-separated organelles such as P granules [9], Cajal bodies [10], stress granules [11], nucleoli [12], etc., share the common property of concentrating different elements locally as prerequisite for the specific biological functions. It has been well acknowledged that the driving forces of phase separation are the weak multivalent attractive interactions [13, 14, 15], originating from the featured protein sequences [16, 17, 5]. Recently, extensive studies have established strong connections of the disease-related mutations in promoting the liquid-liquid phase separation with the formation of pathological gel-like and solid-like aggregates [18, 19, 20, 21], underling the important roles of phase separation in regulating the diseases. Thus, it is imperative to delineate the fundamental physical principles of phase separation, the underlying interactions and the external factors impacting this process.

Environmental factors, such as temperature, pH, salt concentration, etc., can significantly impact the macroscopic properties of phase separation [22, 5]. Notably, IDPs as major players in phase separation, possess an arresting functional advantage that is their high resistance to environmental perturbations [23]. This resilience is largely attributed to the fact that the biological functions of IDPs, realized by the unstructured chains, are more likely to remain consistent under various conditions [24]. Recent experiments have revealed that IDPs in liquid-like droplets retain high degrees of conformational disorders [25, 26]. In particular, Majumdar et al. developed an excimer emission fluorescence technique focusing on intra-molecular distance measurements and identified large-scale conformational expansion during the transition from the the dilute phase to the dense phase for the Alzheimer’s disease-related tau protein [27]. These findings give rise to the hypothesis that the structures of highly extended IDPs in the condensates may also be insensitive to environmental factors. Consequently, the seemingly contradictory environment-(in)dependent macroscopic properties of the condensates and microscopic structures of proteins within the condensates pose an intriguing avenue for further investigation.

Despite IDPs remain conformationally disordered in the phase-separated condensates, the translational diffusions and global motions of the chains are largely repressed in the dense phase with respect to the ones in bulk. This is due to the presence of highly concentrated biomolecules, which lead to high viscosity in the condensates [25, 26, 28, 29]. Recent studies using nuclear magnetic resonance (NMR) [30], single-molecule Förster resonance energy transfer (smFRET) [31] and all-atom molecular dynamics (MD) simulations [31, 32], all demonstrated that IDPs are highly mobile in the condensates and the local chain motions are exceedingly dynamic, associated with rapid breaking and formation of different types of molecular interactions [33]. This highly dynamic chain property induced by the weakly formed, transient interactions, can contribute to the fast exchanges of the binding partners, thus facilitating the efficient realization of biomolecular function in the condensates [34, 35]. Considering the fact that liquid-like chain dynamics within the dense phase are essential for the biological function [36, 7, 37], there is a delicate balance between the stability of the condensates and dynamics of the chain, likely modulated by the underlying molecular interactions.

IDPs in general cannot fold into stable structures due to their low hydrophobicity [38]. Bioinformatic analysis revealed that IDPs typically possess high percentages of charged residues in their sequences and most IDPs are polyampholytes [39]. Studies from both theoretical and experimental aspects have demonstrated that IDP sequences with the fully segregated oppositely charged residues can reduce the dimensions of the chains by collapsing [40, 41], which further has significant impacts on the thermodynamic and kinetic properties of the IDP binding [42, 43]. In addition, the charge distribution patterns of the IDP chains play an important role in phase separation, especially the condensate stability [44, 45, 46]. It has been established that there is a strong correlation between the compaction degree of isolated IDP and its propensity to form the phase-separated condensate [22, 44, 47, 45]. This suggests that the bipolarly charged IDPs, which are capable of forming highly compacted chains in isolated states, exhibit high stability of the phase-separated condensates [44]. As IDPs undergo conformational expansions from collapsed structures at the free state to extended structures in the condensates during phase separation [27], unraveling of highly collapsed IDP chains may contribute to slowing down the process of phase separation. A similar phenomenon was previously observed in the folding-coupled-binding of the bipolarly charged IDP Chz1 to its target histone proteins, where the collapsed IDP chains were deemed as trapping states during the association process [48, 49, 50]. However, the interplay between the stability of the condensates and the kinetic rates of forming condensates in phase separation remains largely elusive.

To address these challenges, we performed MD simulations with a simplified coarse-grained model to investigate the thermodynamics, dynamics and kinetics of phase separation induced by a bipolarly charged IDP system. We observed that the extended IDP conformational ensemble in the condensates are highly conserved with respect to the changes in the interaction strengths and environmental conditions. The global reconfiguration timescales of the IDP chains were found to be slowed down upon condensate formation, while the local chain flexibility remained quite similar in both dilute and dense phases. Notably, we uncovered a non-monotonic relationship between the stability and formation rates of condensates, highlighting the delicate balance between the speed and stability in phase separation. Our study with a focus on exploring the interplay among stability, dynamics and kinetics in phase separation, offers valuable insights into the mechanistic understanding of this complex phenomenon.

## 2 Materials and Methods

### 2.1 Coarse-grained model

Despite recent progress in all-atom MD simulations on identifying different types of interactions stabilizing the dense phase [51, 32, 31], coarse-grained models offer unique advantages in reaching the necessary time scales for studying phase separation processes [52]. Various coarse-grained models have been developed, leading to significant achievements in understanding these slow and complex processes [47, 45, 53, 54, 55, 56]. Here, we employed a simplified coarse-grained model to characterize the thermodynamic stability, chain conformation, dynamic characteristics and kinetic rates of phase separation.

We used a *C*_*α*_ -based coarse-grained model, in which each residue is represented by one bead [57]. Consecutive beads are connected by a harmonic bond with the equilibrium distance set at the typical distance between *C*_*α*_ atoms of two adjacent residues in proteins (*σ* = 0.38 *nm*). Non-bonded interactions between residues separated by more than one bond are described by the Lennard-Jones (LJ) potential and Debye-Hückel potential. The former reflects the short-range pairwise attractive interactions, while the latter accounts for the long-range electrostatic interactions influenced by salt concentrations. The LJ potential (*V*_*LJ*_) is expressed as:

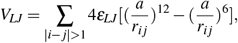

where *r*_*i j*_ is the distance between bead *i* and *j*, the parameter *a* was set to be 0.60 *nm*, mimicking the average diameter of amino acids [58], and *ε*_*LJ*_ controls the depth of the LJ potential well.

The Debye-Hückel model is expressed by:

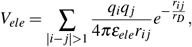

where *q*_*i*_ is the charge of the bead *i, ε*_*ele*_ is the dielectric constant, which was set to be 80 throughout the simulations, and *r*_*D*_ is the Debye screening length, which is dependent on the salt concentration (*C*_*salt*_). The expression of *r*_*D*_ is given by 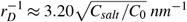, where *C*_0_ is the reference concentration of 1.00 *M* [59, 60, 61]. Thus, higher (lower) salt concentrations lead to stronger (weaker) screening effects of ions on electrostatic interactions, resulting in smaller (biger) values of *r*_*D*_.

We studied the phase separation of an ideal polyampholytic IDP system, which is comprised of 25 lysine (K, positively charged) and 25 glutamic acid (E, negatively charged) residues in one single chain [41, 45]. Specifically, we focused on the system, where the oppositely charged residues are fully segregated in the linear sequence (Figure 1A). This IDP system was previously found to be highly compacted in the dilute phase, exhibiting a high propensity for phase separation [44, 45].

**Figure 1:**
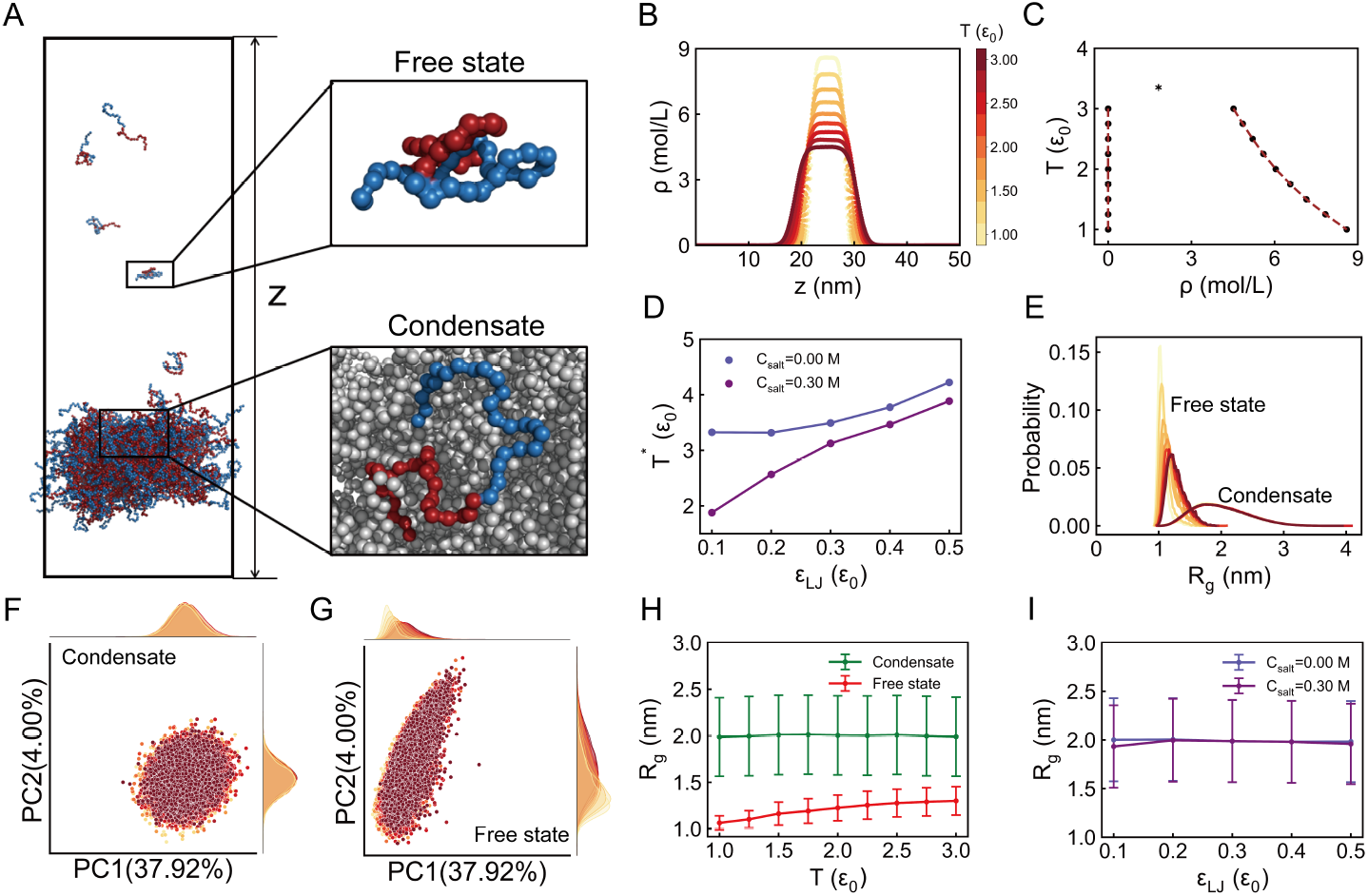
Thermodynamic and conformational properties of IDPs in phase separation. (A) Snapshot of the system after the slab simulation reached equilibration. Zoom-ins are representative structures of IDPs in the dilute phase and dense phase. Positively and negatively charged residues in IDPs are colored in blue and red, respectively. (B) Density profiles along the z-axis varied by temperatures. (C) Phase diagram obtained from the density profile. The critical temperature is denoted as *T* ^∗^. (D) Changes in *T* ^∗^ as a function of interaction strengths (*ε*_*LJ*_) at both high and low salt concentrations (*C*_*salt*_). (E) Distribution of the radius of gyration (*R*_*g*_) for IDPs at the free state and in the condensates varied by temperatures, respectively. (F, G) Two-dimensional principal component analysis (PCA) of the conformation ensembles of IDPs at the free state and in the condensates, respectively. (H) Average *R*_*g*_ of the IDP as a function of temperature at the free state and in the condensates, respectively. (I) Average *R*_*g*_ of the IDP in the condensates as a function of *ε*_*LJ*_ at both high and low *C*_*salt*_, respectively. If not explicitly specified, interaction strength and salt concentration were set to be *ε*_*LJ*_ =0.20 *ε*_0_ and *C*_*salt*_ =0.00 M, respectively. Conformational analyses of IDPs were conducted at *T* = 2.00 *ε*_0_.

### 2.2 Thermodynamic simulations: slab method

Even with the coarse-grained model, it is practically challenging to obtain well-converged thermodynamic properties for the coexistence of two phases in simulations, starting from the fully dispersed and randomly placed configurations. To address this issue, we employed the slab method, which was previously developed by Dignon et al. [53]. The slab simulations were initialized with a slab-like configuration, in which the chains are densely packed in a confined space. This setup can enhance the convergence of the thermodynamic simulation, facilitating the accurate study of phase separation. The slab method has been widely used to study various phase separation processes induced by different types of biomolecules using various models [53, 45, 62], demonstrating the validity of this method. Here, we applied the slab method to conduct the simulations, aiming at investigating the thermodynamic properties of phase separation and conformational characteristics of the IDP chains.

The slab simulations were implemented as follows. Initially, 200 IDP chains, each comprising 50 amino acids, were placed within a square box. Then, an NPT simulation was conducted using Berendsen pressure coupling, with the reference pressure set to 1000 bar in the xyz direction [63]. Through this, we gradually decreased the box size during the simulation till the amino acid concentration reached approximately 3.00 *M*, which is a typical concentration observed in the condensates, experimentally (2.00 to 5.00 *M*) [58]. Then, the box size was fixed with the dense phase appearing in a slab-like configuration. Finally, to achieve the coexistence of the dilute and condensed phases, the box was stretched in the z-direction to 50 *nm*, and NVT simulations were performed (Figure 1A). Analyses were conducted after the system reached equilibration during the NVT simulations.

### 2.3 Kinetic simulations

To investigate the rates of forming the condensates during the phase separation process, we performed kinetic simulations, using the same stretched box employed in the thermodynamic slab simulations. Initial configurations of the kinetic simulations were generated through the thermodynamic simulations performed at high temperature, where only one homogeneous phase was present in the system. For each LJ interaction strength or environmental condition conducive to phase separation, we performed 10 independent simulations, each initialized from a different configuration.

### 2.4 Simulation protocols

We used Gromacs (version 4.5.7) software to perform the MD simulations [64]. Reduced units were used throughout the simulations, except that the length is in the unit of *nm*. Temperature is in the energy unit (*ε*_0_) by multiplying the Boltzmann constant. Time step was set to be 0.0005 *τ*_0_, where *τ*_0_ is the unit of the time. We applied Langevin dynamics with the friction coefficient set as 1.0 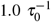. The non-bonded interactions were cut off at the distance of 6*a* = 3.60 *nm*, and the periodic boundary conditions were used in all three dimensions. To explore the effects of interactions and environmental conditions on phase separation, we modulated the strength of the LJ potential (*ε*_*LJ*_), temperature (*T*) and salt concentrations (*C*_*salt*_).

In each thermodynamic simulation, the equilibration step was set to be 5×10^4^ *τ*_0_, which is sufficient for achieving convergence. Kinetic simulations were performed at *T* = 2.00 *ε*_0_, where phase separation could occur for all the interaction strengths and environmental conditions of interest. The kinetic trajectories from the first 500 *τ*_0_ were collected for analysis.

## 3 Results

### 3.1 Conserved conformation ensembles of IDPs in condensates

We investigated the phase separation of an ideal polyampholytic IDP system, consisting of 25 positively and 25 negatively charged residues with a bipolarly charged distribution in the linear sequence. This IDP, corresponding to the KE sequences studied elsewhere [41], exhibits a diblock sequence with the maximum opposite charge segregation (Figure 1A). Previous studies have elucidated the critical roles of electrostatic interactions in inducing single-chain compaction and multiple-chain phase separation in this IDP system [41, 44, 45]. Using the slab simulation method, we identified a coexistence of low- and high-density phases for the IDPs under moderate environmental conditions and the IDPs appeared to be conformationally different between the dilute and dense phases (Figure 1A and 1B). As the temperature increases, the trend of the coexistence of the two phases weakens (Figure 1C). The critical temperature (*T* ^∗^), representing the highest temperature for phase separation to occur, was calculated from the coexistence densities, as detailed in previous studies and SI text [47, 53].

We found that the phase separation of this IDP system can occur at a wide range of the non-bonded LJ interaction strengths (*ε*_*LJ*_) and salt concentrations (*C*_*salt*_) (Figure S3-S12). This suggests that the bipolarly charged IDP naturally possesses a sequence with a strong propensity for phase separation [53]. To investigate the effects of interactions and salt concentrations on phase separation, we calculated the critical temperature (*T* ^∗^) as a function of *ε*_*LJ*_ at both high and low values of *C*_*salt*_ (Figure 1D). The critical temperature characterizes the tendency for IDPs to undergo phase separation, serving as an potential indicator for the stability of the condensates formed in the coexistence phases [53]. We observed monotonically positive correlations of *T* ^∗^ with *ε*_*LJ*_ at both *C*_*salt*_. A comparison of *T* ^∗^ between these two salt concentrations revealed that strong electrostatic interactions favor phase separation. Our results suggest that strengthening the interactions, either through short-range LJ interactions or long-range electrostatic interactions, is capable of enhancing the stability for phase separation of IDPs. Previous research has established a power-law correlation between the degree of single-chain compaction, described by the average radius of gyration (*R*_*g*_) and the stability of multi-chain phase separation (*T* ^∗^) [44, 47, 65]. In our simulations, we observed that *ε*_*LJ*_ and *C*_*salt*_ have significant impacts on the single-chain compaction, further leading to strong correlations between *R*_*g*_ and *T* ^∗^ (Figure S2).

Interestingly, despite the apparent temperature-dependent compaction of IDPs at the free state, we observed that the distributions of *R*_*g*_ for IDPs in the condensates during phase separation remain consistent across temperatures (Figure 1E). Further analysis using principal component analysis (PCA), which considers the spatial distribution of residues relative to the center of mass of the IDPs, also exhibited similar behaviors (Figure 1F and 1G). Additionally, we noted a positive correlation of average *R*_*g*_ with temperature for IDPs at the free state, implying that higher (lower) temperatures result in more extended (compacted) structures. In contrast, IDPs in the condensates appeared to be more extended than those in their free state, and these extended structures showed insensitivity to temperature (Figure 1H).

By varying the interaction strengths and salt concentrations, we observed that the average *R*_*g*_ remained nearly constant for IDPs in the condensates (see Figure 1I). These results suggest that the conformational ensemble of IDPs in condensates is highly conserved and does not change with variations in interactions or environmental conditions. We further conducted a thorough analysis of intra- and inter-chain contacts for IDPs at the free state and in the condensates (Figure S5). We observed a decreasing number of contacts within the chain itself in the free state with increasing temperature, providing an explanation for why IDPs tend to adopt more extended conformations at higher temperatures. In contrast, the number of intrachain contacts for IDPs in the condensates remained nearly constant across different temperatures and environmental conditions. Furthermore, IDPs in the condensates exhibited fewer intra-chain contacts than those at the free state. These results suggest that IDPs are more extended in the condensates, in line with previous simulation and experimental studies [66, 27].

Notably, the number of inter-chain contacts for IDPs in the condensates decreases significantly with increasing temperature (Figure S5). Additionally, changing the interaction strengths and salt concentrations has a significant impact on altering the inter-chain contacts of IDPs in the condensates (Figure S3-S12). Our findings revealed that the conformation ensembles of IDPs remain largely unaltered with respect to changes in interaction strengths, temperatures and salt concentrations (Figure S3-S12). Therefore, the interaction and environmental factors modulate the macroscopic thermodynamic properties of phase separation primarily through changing the microscopic underlying inter-chain interactions of IDPs, which possess the highly conserved, albeit extended, conformational ensembles in the condensates.

### 3.2 Fast dynamics of IDPs in condensates

To assess the liquid-like nature of the dense phase, we calculated the mean squared displacement (MSD) of the IDPs in the condenstates moving along the z-direction as a function of time (Figure 2A). The linear region in the plot suggests that the dynamics of IDPs in the condensates are liquid-like, indicating fluid behavior rather than tightly formed aggregation. Additionally, we observed that IDPs diffuse much faster at the free state than in the condensates. This observation, reminiscent of numerous previous studies [67, 68, 69], is primarily attributed to the high viscosity present in the condensate, resulting from the multivalent interaction network induced by multi-chain of IDPs [14].

**Figure 2:**
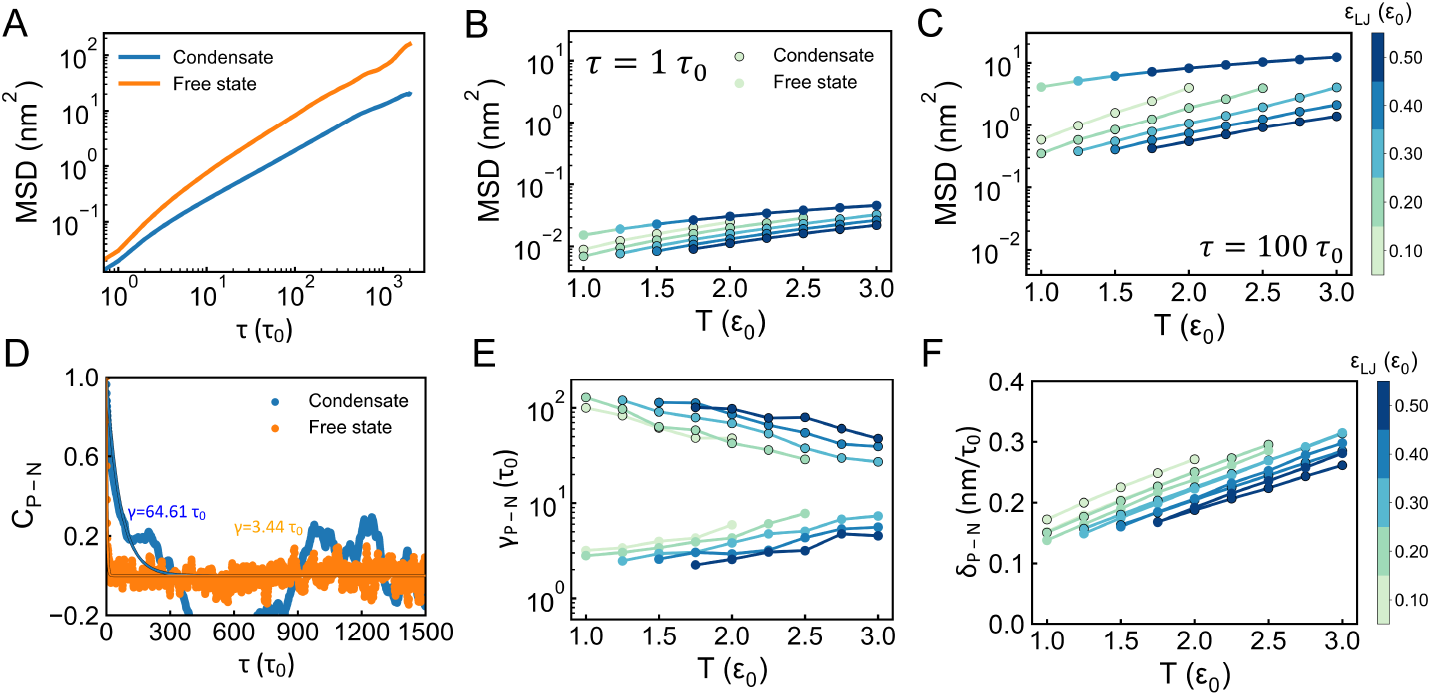
Dynamics of IDPs in the condensates at high salt concentration (*C*_*salt*_ = 0.30 M). (A) Mean squared displacements (MSD) of IDPs at the free state and in the condensates. (B, C) MSD values at the free state and in the condensates with two time intervals *τ* = 1 *τ*_0_ and 100 *τ*_0_. The dots without the black edges represent the data at the free state and those with edges represent the data in the condensates. The same drawing scheme was applied to the following sub-figures (E) and (F). (D) Autocorrelation of the distance between the positively and negatively charged centers (*d*_*P*−*N*_) in the IDP chain as a function of time. The relaxation time *γ*_*P*−*N*_ was obtained by fitting the curve to the single-exponential function (see SI text). (E) Temperature-dependence of *γ*_*P*−*N*_ as a function of *ε*_*LJ*_ at the free state and in the condensates. (F) Rate for describing the IDP conformational dynamics measured by *δ*_*P*−*N*_, which is the change of *d*_*P*−*N*_ per time unit. The results shown in (A) and (D) were obtained with interaction strength and temperature set to be *ε*_*LJ*_ = 0.20 *ε*_0_ and *T* = 2.00 *ε*_0_, respectively.

Recent accumulating evidence has shown that proteins in phase-separated condensates may exhibit different behaviors at various timescales [70, 31, 6, 71, 30]. We extracted the MSD values at different timescales and examined their correlations with interaction strengths and temperatures (Figure 2B and 2C). The gaps between the MSD values at the free state and in the condensates become more significant as the observing timescales increase, indicating that the slowing effects of condensate formation on the translational diffusion of IDPs are more pronounced at longer timescales. Since the diffusion of the IDP at the free state is directly proportional to temperature, the slope of the MSD–T curve indicates the positive temperature effect on facilitating the diffusivity of the chain. Interestingly, we found that although the MSD values are smaller in the condensates than at the free state, the slopes of MSD–T curves are generally steeper, suggesting that temperature has a more profound effect on accelerating the motions of the chain in the condensates. This observation is likely due to the fact that increasing temperature not only increases the kinetic energy of the beads but also loosens the network of the interchain interactions.

We calculated the timescales of the IDP conformational dynamics, represented by the relaxation time *γ* of the autocorrelation function *C*_*P*−*N*_, which focuses on the change in distance between positively and negatively charged centers of the IDP chain (Figure 2D). Our findings indicated that the conformational reconfigurations of IDPs are much slower in the condensates compared to those at the free state. This suggests that formation of the condensates can slow down both translational motions and conformational adaptations of the IDPs, as widely observed in previous studies [30, 31].

To investigate how interaction strengths and temperature influence the timescales of the chain dynamics, we calculated the relaxation time (*γ*_*P*−*N*_) for IDPs to conformationally rearrange their positively and negatively charged centers, at the free state and in the condensates with different temperatures and *ε*_*LJ*_. Strikingly, we observed distinct temperature- and interaction-dependent behaviors of *γ*_*P*−*N*_ for IDPs at the free state and in the condensates (Figure 2E). When the IDP is isolated, weakening the intra-chain interactions induces a more extended and flexible chain, thus slowing down the conformational dynamics. Although increasing temperature accelerates the motions of the beads in the IDP chain, the overall slow chain dynamics led by more extended and flexible conformations can compensate for the gained kinetic energy from the high temperatures, eventually resulting in a negative correlation between temperature and reconfiguration timescales for the IDPs at the free state. In contrast, as demonstrated in Figure 1, strengthening the interactions does not alter the intra-chain conformational ensembles but increases the number of inter-chain contacts in the condenstates. In this regard, stronger (weaker) interactions lead to a more (less) dense phase, resulting in slower (faster) conformational dynamics of IDPs in the condensates. Interchain interactions inside condensates has been deemed as an important factor to slowing of chain dynamics in addition to the viscosity led by the density increase in the dense phase [26]. Furthermore, increasing temperature weakens the interchain interactions associated with highly conserved intra-chain conformations, thereby facilitating the IDP conformational dynamics in the condensates.

We further investigated the short-time chain dynamics by calculating the changes in *d*_*P*−*N*_ per time unit (*δ*_*P*−*N*_, Figure 2F). *δ*_*P*−*N*_ measures the amplitudes of chain conformatonal sampling, thus it reflects the chain flexibility. Different from the reconfiguration timescales of the global chain dynamics, we observed slight enhancements in the chain flexibilty in the condensates with respect to those at the free state. Further-more, temperature has positive effects on enhancing chain flexibility both at the free state and in the condensates. IDPs in the condensates experience high macroscopic viscosity, leading to slowing down the global conformational dynamics of the chain. However, our results suggest that the amplitudes of the chain dynamics at short timescales appear to be largely preserved, similar to those at the free state. Our results echoes previous experiments, where IDPs can maintain their chain flexibility upon phase separation or in the presence of intracellular crowders [72, 73].

### 3.3 Speed-stability balance of phase separation

To study the rates of forming phase-separated condensates, we performed kinetic simulations starting from configurations where only one homogeneous phase exists. We introduced the fraction of inter-chain contacts (*Q*_*atoms*_) of the IDPs formed in the stable condensates to describe the progress of phase separation. *Q*_*atoms*_ is expressed as:

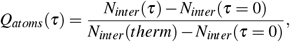

where *N*_*inter*_ is the average number of inter-chain contacts and *N*_*inter*_(*therm*) is *N*_*inter*_ calculated at the stable condensates from the corresponding thermodynamic slab simulations.

We observed a non-linear behavior of *Q*_*atoms*_ evolving with time, reminiscent of the multi-step nucleation growth and assembly process in phase separation observed in recent experiments [74, 75] (Figure 3A and 3B), at each interaction strength. Specifically, *Q*_*atoms*_ increases abruptly at the beginning of the phase separation process, indicating the rapid formation of a significant number of inter-chain contacts in the early stages. The subsequent establishment of inter-chain contacts appears to be relatively slow, possibly involving the droplet-coalesced condensate-merging process, which can be hindered by the slow thermal diffusivity of the condensates [76, 77, 78]. To compare the kinetic rates of phase separation for different interaction strengths of the IDPs, we extracted the *Q*_*atoms*_ values at the time point *τ* = 300 *τ*_0_ (Figure 3C). We found that strengthening the interactions of IDPs at low salt concentration (*C*_*salt*_ = 0.00 *M*) accelerates the formation of condensates. However, the effects of the interaction strength of IDPs on the kinetics of phase separation are non-monotonic at high salt concentration (*C*_*salt*_ = 0.30 *M*). In particular, the rate of forming the phase-separated condensates is the slowest when *ε*_*LJ*_ = 0.20 *ε*_0_. A similar observation can be made when the time point was chosen to be 400 *τ*_0_ (Figure S14).

**Figure 3:**
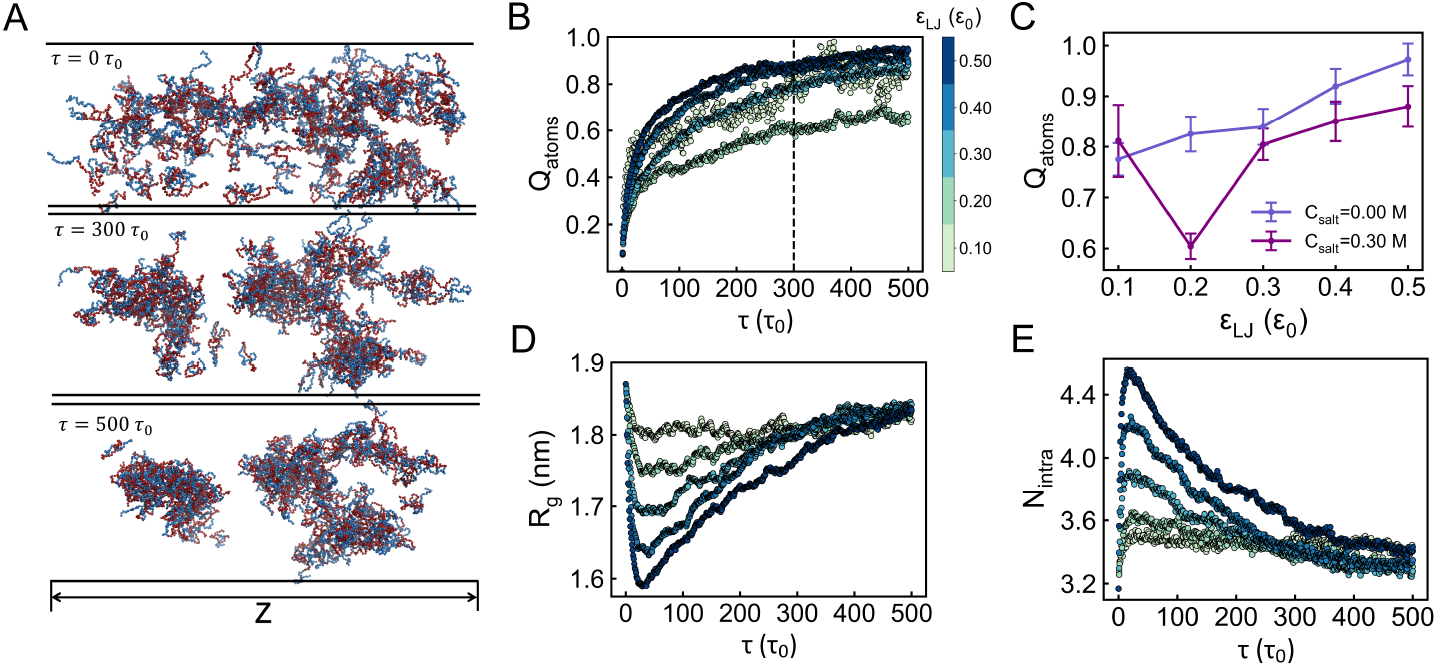
Kinetics of phase separation at high salt concentration (*C*_*salt*_ = 0.30 *M*). The temperature was set to be *T* = 2.00 *ε*_0_, where phase separation can occur steadily at various interaction strengths, as demonstrated in thermodynamic slab simulations. (A) Snapshots of the phase separation process at time points *τ* = 0 *τ*_0_, 300 *τ*_0_, and 500 *τ*_0_. The structures were extracted from the simulation with interaction strength set to be *ε*_*LJ*_ = 0.20 *ε*_0_. (B) Evolution of phase separation along with time. The process is described by *Q*_*atoms*_, which is the fraction of inter-chain contacts formed in the thermodynamic slab simulations. (C) *Q*_*atoms*_ values at the time *τ* = 300 *τ*_0_. (D) Evolution of the average *R*_*g*_ in the process of phase separation. (E) Evolution of the average number of intra-chain contacts (*N*_*intra*_) in the process of phase separation.

Multivalent interactions occurring at the inter-chain scale have been deemed important in facilitating the phase separation of IDPs [14]. Stronger (weaker) inter-chain interactions should lead to faster (slower) formation of condensates in phase separation. Consequently, the aberrant non-monotonic relationship between the kinetic rates and the interaction strengths may be attributed to intra-chain interactions. Hence, we monitored the evolution of intra-chain conformational properties, described by the average *R*_*g*_ of the chain and the average number of intra-chain contacts (*N*_*intra*_) in the IDP, during phase separation (Figure 3D and 3E). We observed abrupt chain compaction associated with significant establishment of intra-chain contacts in IDPs at the early stages of the phase separation processes. These observations are more pronounced with a larger value of *ε*_*LJ*_, where IDPs exhibit more conformational collapse when they start to phase separate, compared to those in the condensates. As demonstrated in our thermodynamic analyses (Figure 1), the conformational ensembles of IDPs are highly conserved in the condensates with different interaction strengths. Thus, forming the extended IDP conformations in the condensates inevitably leads to opening up of the chains during phase separation. Strong interactions of IDPs may contribute to slowing down the processs, particularly when *ε*_*LJ*_ is large enough to induce highly compacted IDP conformations at the free state. On the other hand, strong interactions of IDPs lead to the high degrees of chain compaction at the free state, further resulting in high thermo-dynamic stability for phase separation. In this regard, there is a balance between kinetic efficiency and thermodynamic stability of the forming the condensates, delicately modulated by the chain interactions and external environmental factors. There-fore, we suggest that the high stability of the condenstate does not always imply the fast formation of phase separation and vice versa.

## 4 Discussion and Conclusions

We performed MD simulations using a simplified coarse-grained model to study phase separation of bipolarly charged IDPs. This IDP system, which is an ideal polyampholytic chain, has a fully segregated oppositely charged residues in the linear sequence. Our investigation focused on delineating the thermodynamic stability of the condensates, the conformational properties and chain dynamics of the IDPs in the condensates and the kinetic rate of phase separation under various interaction strengths and environmental conditions.

Consistent with previous studies from theoretical, computational and experimental aspects [22, 44, 47, 45], we also observed that the highly collapsed IDP resulting from strong chain interactions lead to the highly stable phase-separated condensates. This implies a similarity between the inter-chain interactions stabilizing the IDP-rich condensates and the intrachain interactions driving the compaction of IDPs [14, 79, 80]. Interestingly, different interaction strengths led to different degrees of IDP chain compaction but highly conserved, extended IDP conformational ensembles in the condensates (Figure 4). Further alterations in environmental conditions do not change the conformational ensembles of the IDPs, suggesting that formation of the condensates may serve as a protection shield, wherein IDPs can maintain their molecular-scale conformation ensembles. As the structural properties are essential for exerting the correct biological function, our results suggest that for-mation the condensates is beneficial to proteins from the functional perspective, as they are more resistant to the alterations of external stimuli than at the free state.

**Figure 4:**
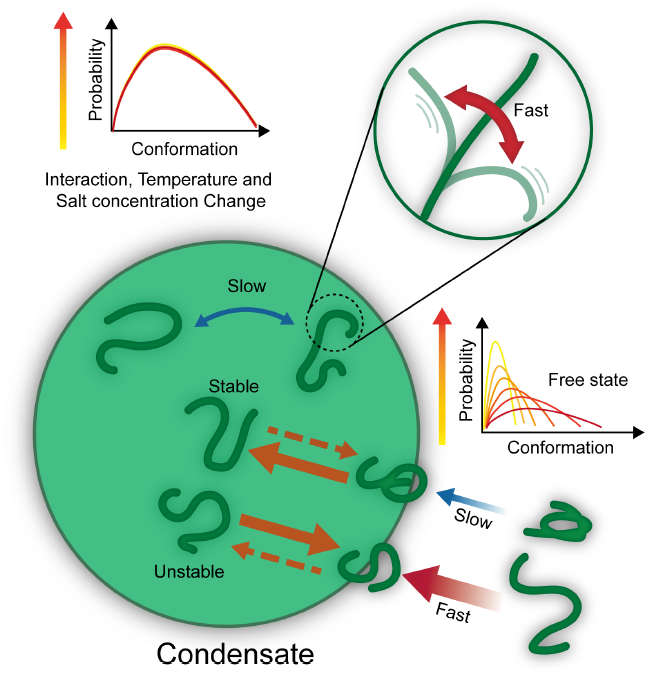
Schematic illustration summarizing the key findings of this work. Different interactions and environmental conditions result in different stabilities of phase-separated condensates. In contrast to the free state, the conformational ensembles of IDPs in the condensates appear to be highly conserved. Despite the translational diffusion and conformational dynamics of IDPs being slowed in the condensates, rapid chain dynamics at short timescales remain comparable to those in the free state. The compacted nature of IDPs in the free state, while beneficial for condensate stability, necessitates chain opening during phase separation, thereby contributing to slowing down the kinetics of phase separation.

Due to the high viscosity caused by multi-chain assembly, IDPs in the dense phase have been consistently characterized to exhibit the slowing down of translational diffusion and conformational dynamics, compared to when they are in the dilute phase [67, 68, 69]. Here, we observed distinct interaction- and temperature-dependent behaviors on the reconfiguration timescales of global chain dynamics for IDPs at the free state and in the condensates. This is likely due to the fact that changing interactions or environmental conditions alters the network of multi-chain interactions, further impacting the motions and chain dynamics of IDPs in the condensates. At the free state, weak interactions and high temperature induce broad conformational distributions of IDPs, thus slowing down the reconfiguration dynamics of the global chain conformation. In contrast, due to the protective effects provided by the condensates, the conformational distributions of IDPs remain similar in dense phase under different conditions, while the reconfiguration chain dynamics of IDPs are accelerated by the weak inter-chain interactions led by the small *ε*_*LJ*_ and high temperature. Our findings highlight that the macroscopic thermody-namic properties of the condensates and global chain dynamics of the IDPs in the dense phase are intimately modulated by multiple factors, primarily through inter-chain interactions [71]. We also observed that the fast chain sampling dynamics of isolated IDPs at short timescales are preserved in the condensates (Figure 4). These findings are in good agreement with a recent study that the rapid breaking and forming of inter-chain contacts in the condensates were detected through a combination of single-molecule spectroscopy and all-atom MD simulations [31].

We observed a non-monotonic correlation between the interaction strengths of the chains and the kinetic rates for forming the condensates (Figure 4). This result underscores the subtle balance between the speed and stability in biomolecular phase separation. It is widely recognized that inter-chain interactions play a critical role in phase separation, as they must be strong enough for the formation of stable condensates and weak enough to ensure the liquid-like dynamics [36, 7]. Our kinetic study emphasizes the importance of the moderate strengths of chain interactions in promoting the formation of phase-separated condensates. Taken together, phase separation is optimized by multiple factors through underlying molecular interactions [14], ensuring the efficient formation of stable condensates, wherein the rapid liquid-like chain dynamics involving interaction forming and breaking can occur, simultaneously.

In summary, our work uncovered the role of phase-separated condensates in preserving the conformational properties of proteins under various internal and external conditions, serving as a pronounced advantage for maintaining structure-function relationships of proteins in the dense phase [23]. Furthermore, the observed fast short-time chain dynamics against the slow global reconfiguration dynamics at the background in the condensates may contribute to aiding efficient biochemical reactions [36, 7, 37]. We demonstrated that moderate strengths of chain interactions are essential to balance speed and stability for optimizing the phase separation process. Our studies offer molecular-level insights into the interplay of stability, dynamics and kinetics in phase separation, providing useful guidance for engineering biomolecular condensate.

## Acknowledgement

X.C. acknowledges the support from the National Natural Science Foundation of China (Grant No. 32201020), the general program (Grant No. 2023A04J0083) and the Guangzhou-HKUST(GZ) joint funding program (Grant No 2023A03J0060) of the Guangzhou Municipal Science and Technology Project, and Guangdong Scientific Research Platform and Projects for the Higher-educational Institution & Education Science Planning Scheme (Grant No. 2023KTSCX169). X.C. was also partly supported by the Municipal Key Laboratory Construction program of the Guangzhou Municipal Science and Technology Project (Grant No. 2023A03J0003).

## References

[1] Avinash Patel, Hyun O Lee, Louise Jawerth, Shovamayee Maharana, Marcus Jahnel, Marco Y Hein, Stoyno Stoynov, Julia Mahamid, Shambaditya Saha, Titus M Franzmann, et al. A liquid-to-solid phase transition of the als protein fus accelerated by disease mutation. Cell, 162(5):1066–1077, 2015.

[2] Susanne Wegmann, Bahareh Eftekharzadeh, Katharina Tepper, Katarzyna M Zoltowska, Rachel E Bennett, Simon Dujardin, Pawel R Laskowski, Danny MacKenzie, Tarun Kamath, Caitlin Commins, et al. Tau protein liquid–liquid phase separation can initiate tau aggregation. The EMBO journal, 37(7):e98049, 2018.

[3] Nicholas M Kanaan, Chelsey Hamel, Tessa Grabinski, and Benjamin Combs. Liquid-liquid phase separation induces pathogenic tau conformations in vitro. Nature communications, 11(1):2809, 2020.

[4] Soumik Ray, Nitu Singh, Rakesh Kumar, Komal Patel, Satyaprakash Pandey, Debalina Datta, Jaladhar Mahato, Rajlaxmi Panigrahi, Ambuja Navalkar, Surabhi Mehra, et al. α-synuclein aggregation nucleates through liquid–liquid phase separation. Nature chemistry, 12(8):705–716, 2020.

[5] José A Villegas, Meta Heidenreich, and Emmanuel D Levy. Molecular and environmental determinants of biomolecular condensate formation. Nature Chemical Biology, 18(12):1319–1329, 2022.

[6] Bryan A Gibson, Claudia Blaukopf, Tracy Lou, Lifeng Chen, Lynda K Doolittle, Ilya Finkelstein, Geeta J Narlikar, Daniel W Gerlich, and Michael K Rosen. In diverse conditions, intrinsic chromatin condensates have liquid-like material properties. Proceedings of the National Academy of Sciences, 120(18):e2218085120, 2023.

[7] Salman F Banani, Hyun O Lee, Anthony A Hyman, and Michael K Rosen. Biomolecular condensates: organizers of cellular biochemistry. Nature reviews Molecular cell biology, 18(5):285–298, 2017.

[8] Xiaoyu Song, Fengrui Yang, Tongtong Yang, Yong Wang, Mingrui Ding, Linge Li, Panpan Xu, Shuaiyu Liu, Ming Dai, Changbiao Chi, et al. Phase separation of eb1 guides microtubule plusend dynamics. Nature Cell Biology, 25(1):79–91, 2023.

[9] Geraldine Seydoux. The p granules of c. elegans: a genetic model for the study of rna–protein condensates. Journal of molecular biology, 430(23):4702–4710, 2018.

[10] Glenn E Morris. The cajal body. Biochimica et Biophysica Acta (BBA)-Molecular Cell Research, 1783(11):2108–2115, 2008.

[11] David SW Protter and Roy Parker. Principles and properties of stress granules. Trends in cell biology, 26(9):668–679, 2016.

[12] François-Michel Boisvert, Silvana Van Koningsbruggen, Joaquín Navascués, and Angus I Lamond. The multifunctional nucleolus. Nature reviews Molecular cell biology, 8(7):574–585, 2007.

[13] Clifford P Brangwynne, Peter Tompa, and Rohit V Pappu. Polymer physics of intracellular phase transitions. Nature Physics, 11(11):899–904, 2015.

[14] Gregory L Dignon, Robert B Best, and Jeetain Mittal. Biomolecular phase separation: from molecular driving forces to macroscopic properties. Annual review of physical chemistry, 71:53–75, 2020.

[15] Benjamin S Schuster, Gregory L Dignon, Wai Shing Tang, Fleurie M Kelley, Aishwarya Kanchi Ranganath, Craig N Jahnke, Alison G Simpkins, Roshan Mammen Regy, Daniel A Hammer, Matthew C Good, et al. Identifying sequence perturbations to an intrinsically disordered protein that determine its phase-separation behavior. Proceedings of the National Academy of Sciences, 117(21):11421–11431, 2020.

[16] Erik W Martin, Alex S Holehouse, Ivan Peran, Mina Farag, J Jeremias Incicco, Anne Bremer, Christy R Grace, Andrea Soranno, Rohit V Pappu, and Tanja Mittag. Valence and patterning of aromatic residues determine the phase behavior of prion-like domains. Science, 367(6478):694–699, 2020.

[17] Wade Borcherds, Anne Bremer, Madeleine B Borgia, and Tanja Mittag. How do intrinsically disordered protein regions encode a driving force for liquid–liquid phase separation? Current opinion in structural biology, 67:41–50, 2021.

[18] Natalia B Nedelsky and J Paul Taylor. Bridging biophysics and neurology: aberrant phase transitions in neurodegenerative disease. Nature Reviews Neurology, 15(5):272–286, 2019.

[19] Simon Alberti and Dorothee Dormann. Liquid–liquid phase separation in disease. Annual review of genetics, 53:171–194, 2019.

[20] Phuong H Nguyen, Ayyalusamy Ramamoorthy, Bikash R Sahoo, Jie Zheng, Peter Faller, John E Straub, Laura Dominguez, Joan-Emma Shea, Nikolay V Dokholyan, Alfonso De Simone, et al. Amyloid oligomers: A joint experimental/computational perspective on alzheimers disease, parkinsons disease, type ii diabetes, and amyotrophic lateral sclerosis. Chemical reviews, 121(4):2545–2647, 2021.

[21] Yi Zhou, Wennan Chang, Xiaoyu Lu, Jin Wang, Chi Zhang, and Ying Xu. Acid-base homeostasis and implications to the phenotypic behaviors of cancer. Genomics, Proteomics & Bioinformatics, 2022.

[22] Joshua A Riback, Christopher D Katanski, Jamie L Kear-Scott, Evgeny V Pilipenko, Alexandra E Rojek, Tobin R Sosnick, and D Allan Drummond. Stress-triggered phase separation is an adaptive, evolutionarily tuned response. Cell, 168(6):1028–1040, 2017.

[23] Zhirong Liu and Yongqi Huang. Advantages of proteins being disordered. Protein Science, 23(5):539–550, 2014.

[24] Vladimir N Uversky. Unusual biophysics of intrinsically disordered proteins. Biochimica et Biophysica Acta (BBA)-Proteins and Proteomics, 1834(5):932–951, 2013.

[25] Sean E Reichheld, Lisa D Muiznieks, Fred W Keeley, and Simon Sharpe. Direct observation of structure and dynamics during phase separation of an elastomeric protein. Proceedings of the National Academy of Sciences, 114(22):E4408–E4415, 2017.

[26] Jacob P Brady, Patrick J Farber, Ashok Sekhar, Yi-Hsuan Lin, Rui Huang, Alaji Bah, Timothy J Nott, Hue Sun Chan, Andrew J Baldwin, Julie D Forman-Kay, et al. Structural and hydrodynamic properties of an intrinsically disordered region of a germ cell-specific protein on phase separation. Proceedings of the National Academy of Sciences, 114(39):E8194–E8203, 2017.

[27] Anupa Majumdar, Priyanka Dogra, Shiny Maity, and Samrat Mukhopadhyay. Liquid–liquid phase separation is driven by large-scale conformational unwinding and fluctuations of intrinsically disordered protein molecules. The journal of physical chemistry letters, 10(14):3929–3936, 2019.

[28] Anastasia C Murthy, Gregory L Dignon, Yelena Kan, Gül H Zerze, Sapun H Parekh, Jeetain Mittal, and Nicolas L Fawzi. Molecular interactions underlying liquidliquid phase separa-tion of the fus low-complexity domain. Nature structural & molecular biology, 26(7):637–648, 2019.

[29] Tae Hun Kim, Brian Tsang, Robert M Vernon, Nahum Sonenberg, Lewis E Kay, and Julie D Forman-Kay. Phosphodependent phase separation of fmrp and caprin1 recapitulates regulation of translation and deadenylation. Science, 365(6455):825–829, 2019.

[30] Serafima Guseva, Vincent Schnapka, Wiktor Adamski, Damien Maurin, Rob WH Ruigrok, Nicola Salvi, and Martin Blackledge. Liquid–liquid phase separation modifies the dynamic properties of intrinsically disordered proteins. Journal of the American Chemical Society, 145(19):10548–10563, 2023.

[31] Nicola Galvanetto, Milos? T Ivanovic, Aritra Chowdhury, Andrea Sottini, Mark F Nüesch, Daniel Nettels, Robert B Best, and Benjamin Schuler. Extreme dynamics in a biomolecular condensate. Nature, 619(7971):876–883, 2023.

[32] Wenwei Zheng, Gregory L Dignon, Nina Jovic, Xichen Xu, Roshan M Regy, Nicolas L Fawzi, Young C Kim, Robert B Best, and Jeetain Mittal. Molecular details of protein condensates probed by microsecond long atomistic simulations. The Journal of Physical Chemistry B, 124(51):11671–11679, 2020.

[33] Nicolas L Fawzi, Sapun H Parekh, and Jeetain Mittal. Biophysical studies of phase separation integrating experimental and computational methods. Current Opinion in Structural Biology, 70:78–86, 2021.

[34] Andrea Sottini, Alessandro Borgia, Madeleine B Borgia, Katrine Bugge, Daniel Nettels, Aritra Chowdhury, Pétur O Heidarsson, Franziska Zosel, Robert B Best, Birthe B Kragelund, et al. Polyelectrolyte interactions enable rapid association and dissociation in high-affinity disordered protein complexes. Nature communications, 11(1):5736, 2020.

[35] Pétur O Heidarsson, Davide Mercadante, Andrea Sottini, Daniel Nettels, Madeleine B Borgia, Alessandro Borgia, Sinan Kilic, Beat Fierz, Robert B Best, and Benjamin Schuler. Release of linker histone from the nucleosome driven by polyelectrolyte competition with a disordered protein. Nature Chemistry, 14(2):224–231, 2022.

[36] Yongdae Shin and Clifford P Brangwynne. Liquid phase condensation in cell physiology and disease. Science, 357(6357):eaaf4382, 2017.

[37] Christopher D Reinkemeier and Edward A Lemke. Synthetic biomolecular condensates to engineer eukaryotic cells. Current Opinion in Chemical Biology, 64:174–181, 2021.

[38] Vladimir N Uversky. What does it mean to be natively unfolded? European journal of biochemistry, 269(1):2–12, 2002.

[39] Megan Sickmeier, Justin A Hamilton, Tanguy LeGall, Vladimir Vacic, Marc S Cortese, Agnes Tantos, Beata Szabo, Peter Tompa, Jake Chen, Vladimir N Uversky, et al. Disprot: the database of disordered proteins. Nucleic acids research, 35(suppl 1):D786–D793, 2007.

[40] Sonja Müller-Späth, Andrea Soranno, Verena Hirschfeld, Hagen Hofmann, Stefan Rüegger, Luc Reymond, Daniel Nettels, and Benjamin Schuler. Charge interactions can dominate the dimensions of intrinsically disordered proteins. Proceedings of the National Academy of Sciences, 107(33):14609–14614, 2010.

[41] Rahul K Das and Rohit V Pappu. Conformations of intrinsically disordered proteins are influenced by linear sequence distributions of oppositely charged residues. Proceedings of the National Academy of Sciences, 110(33):13392–13397, 2013.

[42] Samrat Mukhopadhyay, Rajaraman Krishnan, Edward A Lemke, Susan Lindquist, and Ashok A Deniz. A natively unfolded yeast prion monomer adopts an ensemble of collapsed and rapidly fluctuating structures. Proceedings of the National Academy of Sciences, 104(8):2649–2654, 2007.

[43] Albert H Mao, Scott L Crick, Andreas Vitalis, Caitlin L Chicoine, and Rohit V Pappu. Net charge per residue modulates conformational ensembles of intrinsically disordered proteins. Proceedings of the National Academy of Sciences, 107(18):8183–8188, 2010.

[44] Yi-Hsuan Lin and Hue Sun Chan. Phase separation and singlechain compactness of charged disordered proteins are strongly correlated. Biophysical Journal, 112(10):2043–2046, 2017.

[45] Suman Das, Alan N Amin, Yi-Hsuan Lin, and Hue Sun Chan. Coarse-grained residue-based models of disordered protein condensates: utility and limitations of simple charge pattern parameters. Physical Chemistry Chemical Physics, 20(45):28558–28574, 2018.

[46] Wen-Ting Chu and Jin Wang. Thermodynamic and sequential characteristics of phase separation and droplet formation for an intrinsically disordered region/protein ensemble. PLoS computational biology, 17(3):e1008672, 2021.

[47] Gregory L Dignon, Wenwei Zheng, Robert B Best, Young C Kim, and Jeetain Mittal. Relation between single-molecule properties and phase behavior of intrinsically disordered proteins. Proceedings of the National Academy of Sciences, 115(40):9929–9934, 2018.

[48] Xiakun Chu, Yong Wang, Linfeng Gan, Yawen Bai, Wei Han, Erkang Wang, and Jin Wang. Importance of electrostatic interactions in the association of intrinsically disordered histone chaperone chz1 and histone h2a. z-h2b. PLoS computational biology, 8(7):e1002608, 2012.

[49] Xiakun Chu, Linfeng Gan, Erkang Wang, and Jin Wang. Quantifying the topography of the intrinsic energy landscape of flexible biomolecular recognition. Proceedings of the National Academy of Sciences, 110(26):E2342–E2351, 2013.

[50] Xiakun Chu, Zucai Suo, and Jin Wang. Investigating the trade-off between folding and function in a multidomain y-family dna polymerase. Elife, 9:e60434, 2020.

[51] Sarah Rauscher and Régis Pomes. The liquid structure of elastin. Elife, 6:e26526, 2017.

[52] Gregory L Dignon, Wenwei Zheng, and Jeetain Mittal. Simulation methods for liquid–liquid phase separation of disordered proteins. Current opinion in chemical engineering, 23:92–98, 2019.

[53] Gregory L Dignon, Wenwei Zheng, Young C Kim, Robert B Best, and Jeetain Mittal. Sequence determinants of protein phase behavior from a coarse-grained model. PLoS computational biology, 14(1):e1005941, 2018.

[54] Stefan Roberts, Tyler S Harmon, Jeffrey L Schaal, Vincent Miao, Kan Li, Andrew Hunt, Yi Wen, Terrence G Oas, Joel H Collier, Rohit V Pappu, et al. Injectable tissue integrating networks from recombinant polypeptides with tunable order. Nature Materials, 17(12):1154–1163, 2018.

[55] Yiming Tang, Santu Bera, Yifei Yao, Jiyuan Zeng, Zenghui Lao, Xuewei Dong, Ehud Gazit, and Guanghong Wei. Prediction and characterization of liquid-liquid phase separation of minimalistic peptides. Cell Reports Physical Science, 2(9), 2021.

[56] Fei Liu and Jin Wang. Atp acts as a hydrotrope to regulate the phase separation of nbdy clusters. JACS Au, 3(9):2578–2585, 2023.

[57] Alexander J Pak and Gregory A Voth. Advances in coarsegrained modeling of macromolecular complexes. Current opinion in structural biology, 52:119–126, 2018.

[58] Roshan Mammen Regy, Wenwei Zheng, and Jeetain Mittal. Using a sequence-specific coarse-grained model for studying protein liquid–liquid phase separation. In Methods in enzymology, volume 646, pages 1–17. 2021.

[59] Ariel Azia and Yaakov Levy. Nonnative electrostatic interactions can modulate protein folding: molecular dynamics with a grain of salt. Journal of molecular biology, 393(2):527–542, 2009.

[60] Xiakun Chu and Jin Wang. Position-, disorder-, and salt-dependent diffusion in binding-coupled-folding of intrinsically disordered proteins. Physical Chemistry Chemical Physics, 21(10):5634–5645, 2019.

[61] Xiakun Chu, Zucai Suo, and Jin Wang. Investigating the conformational dynamics of a y-family dna polymerase during its folding and binding to dna and a nucleotide. JACS Au, 2(2):341–356, 2022.

[62] David De Sancho. Phase separation in amino acid mixtures is governed by composition. Biophysical Journal, 121(21):4119–4127, 2022.

[63] Herman JC Berendsen, JPM van Postma, Wilfred F Van Gunsteren, ARHJ DiNola, and Jan R Haak. Molecular dynamics with coupling to an external bath. The Journal of chemical physics, 81(8):3684–3690, 1984.

[64] David Van Der Spoel, Erik Lindahl, Berk Hess, Gerrit Groenhof, Alan E Mark, and Herman JC Berendsen. Gromacs: fast, flexible, and free. Journal of computational chemistry, 26(16):1701–1718, 2005.

[65] Han-Yi Chou and Aleksei Aksimentiev. Single-protein collapse determines phase equilibria of a biological condensate. The Journal of Physical Chemistry Letters, 11(12):4923–4929, 2020.

[66] Linge Li and Zhonghuai Hou. Crosslink-induced conformation change of intrinsically disordered proteins have a nontrivial effect on phase separation dynamics and thermodynamics. The Journal of Physical Chemistry B, 2023.

[67] Ming-Tzo Wei, Shana Elbaum-Garfinkle, Alex S Holehouse, Carlos Chih-Hsiung Chen, Marina Feric, Craig B Arnold, Rodney D Priestley, Rohit V Pappu, and Clifford P Brangwynne. Phase behaviour of disordered proteins underlying low density and high permeability of liquid organelles. Nature chemistry, 9(11):1118–1125, 2017.

[68] Simon Alberti, Amy Gladfelter, and Tanja Mittag. Considerations and challenges in studying liquid-liquid phase separation and biomolecular condensates. Cell, 176(3):419–434, 2019.

[69] Zheng Wang, Jizhong Lou, and Hong Zhang. Essence determines phenomenon: assaying the material properties of biological condensates. Journal of Biological Chemistry, 298(4), 2022.

[70] Rashik Ahmed and Julie D Forman-Kay. Nmr insights into dynamic, multivalent interactions of intrinsically disordered regions: from discrete complexes to condensates. Essays in Biochemistry, 66(7):863–873, 2022.

[71] Anton Abyzov, Martin Blackledge, and Markus Zweckstetter. Conformational dynamics of intrinsically disordered proteins regulate biomolecular condensate chemistry. Chemical Reviews, 122(6):6719–6748, 2022.

[72] Kathleen A Burke, Abigail M Janke, Christy L Rhine, and Nicolas L Fawzi. Residue-by-residue view of in vitro fus granules that bind the c-terminal domain of rna polymerase ii. Molecular cell, 60(2):231–241, 2015.

[73] Iwo König, Andrea Soranno, Daniel Nettels, and Benjamin Schuler. Impact of in-cell and in-vitro crowding on the conformations and dynamics of an intrinsically disordered protein. Angewandte Chemie, 133(19):10819–10824, 2021.

[74] Erik W Martin, Tyler S Harmon, Jesse B Hopkins, Srinivas Chakravarthy, J Jeremías Incicco, Peter Schuck, Andrea Soranno, and Tanja Mittag. A multi-step nucleation process determines the kinetics of prion-like domain phase separation. Nature communications, 12(1):4513, 2021.

[75] Chengqian Yuan, Qi Li, Ruirui Xing, Junbai Li, and Xuehai Yan. Peptide self-assembly through liquid-liquid phase sepa-ration. Chem, 2023.

[76] Daniel SW Lee, Ned S Wingreen, and Clifford P Brangwynne. Chromatin mechanics dictates subdiffusion and coarsening dynamics of embedded condensates. Biophysical Journal, 120(3):318a, 2021.

[77] Ibraheem Alshareedah, Taranpreet Kaur, and Priya R Banerjee. Methods for characterizing the material properties of biomolecular condensates. In Methods in enzymology, volume 646, pages 143–183. 2021.

[78] Louise Jawerth, Elisabeth Fischer-Friedrich, Suropriya Saha, Jie Wang, Titus Franzmann, Xiaojie Zhang, Jenny Sachweh, Martine Ruer, Mahdiye Ijavi, Shambaditya Saha, et al. Protein condensates as aging maxwell fluids. Science, 370(6522):1317–1323, 2020.

[79] Jeong-Mo Choi, Alex S Holehouse, and Rohit V Pappu. Physical principles underlying the complex biology of intracellular phase transitions. Annual review of biophysics, 49:107–133, 2020.

[80] Xuewei Dong, Santu Bera, Qin Qiao, Yiming Tang, Zenghui Lao, Yin Luo, Ehud Gazit, and Guanghong Wei. Liquid–liquid phase separation of tau protein is encoded at the monomeric level. The journal of physical chemistry letters, 12(10):2576–2586, 2021.

