## Supplementary material for "Balancing stability, dynamics and kinetics in phase separation of intrinsically disordered proteins": SI Text

### Additional Materials and Methods

#### Density profile fitting

To determine the density profile along the z-direction in the thermodynamic slab simulation, we employed the following sigmoidal equation to calculate the concentration of amino acids ($\rho_z$) at a specific coordinate $z$ (Figure S1) [1, 2]:

$$\rho_z = \frac{1}{2}(\rho_H + \rho_L) - \frac{1}{2}(\rho_H - \rho_L)tanh(\frac{z - z_0}{d}),$$

where $\rho_H$ and $\rho_L$ represent concentrations at the dense and dilute phases, respectively. Parameters such as $z_0$ (center position of the dense phase) and $d$ (half-width of the maximum density of the dense phase) were obtained by fitting simulation data to the expression of $\rho_z$ (Figure S1).

Following the determination of $\rho_H$ and $\rho_L$ at various temperatures, the critical temperature $T^*$ was determined by fitting the data to the equation [1, 2]:

$$\rho_H - \rho_L = A(T^* - T)^\beta,$$

where $\beta$ is the critical exponent, which was set to be 0.325, and $A$ is the fitting parameter.

The critical concentration $\rho^*$ was obtained through the following equation:

$$\frac{\rho_H + \rho_L}{2} = \rho^* + B(T^* - T),$$

where $B$ is the fitting parameter and remains constant once $T^*$ is determined.

#### Principal component analysis (PCA)

In addition to $R_g$ and $d_{P-N}$, we performed PCA analysis on the conformational ensembles of the IDP chains at the free state and in the condensates. The PCA plots were generated based on the structural vector $\boldsymbol{D}$, described by the following expression:

$$D_i = |\boldsymbol{r_i} - \boldsymbol{R}|,$$

where $\boldsymbol{r_i}$ represents the Cartesian coordinates of the $i$th residue of the chain, $\boldsymbol{R}$ is the Cartesian coordinate of the center of mass of the chain, and $D_i$ is the $i$th element of the vector $\boldsymbol{D}$. Thus, we used the distance of each bead from the center of mass of the chain as the feature to describe the conformation of the IDP chain. In practice, we used the "sklearn" package of Python to perform PCA analysis.

#### Number of intra-chain and inter-chain contacts

We used the number of spatially proximal beads to quantify the number of contacts formed by the residue of interest. The cut-off distance for counting the contacts was set to be 1 *nm*. We have also made tests on various values of cut-off distance and found no significant changes in our results. The contacts were subsequently classified into the intra-chain and inter-chain contacts.

Direct counting of the contacts formed in the condensates demands significant amount of computational efforts. Therefore, we devised a special purpose-built computational procedure by using the K-Nearest Neighbor (KNN) algorithm [3, 4]. In practice, we collected the coordinates of all residues at each time frame during the simulations. Then, the coordinates of $50 \times 200$ residues were fed into the KNN algorithm, which will automatically calculate the distances between any two residues. Finally, the contact number of each residue was rapidly obtained after setting up the cut-off distance.

#### Mean square displacement (MSD)

To study the diffusive characteristics of the IDP chains at free state and in the condensates. We calculated the MSD of the center of mass of the chain along the z-direction. The MSD was calculated using the following expression:

$$MSD(\tau) = \sum_{t=1}^{t=t_{max}-\tau} \frac{Z(t+\tau) - Z(t)}{t_{max} - \tau},$$

where $Z$ is the z-coordinate of the center of mass of the chain, $t$ denotes the simulation time frame, $t_{max}$ is the maximum simulation time and $\tau$ represents the timescale of interest [5].

#### Autocorrelation function

We used autocorrelation function of $d_{P-N}$ ($C_{P-N}$) to characterize the global conformational dynamics of the IDP chain. The expression of $C_{P-N}$ is given by:

$$C_{P-N}(\tau) = \frac{< (d_{P-N}(\tau) - \overline{d_{P-N}})(d_{P-N}(\tau = 0) - \overline{d_{P-N}}) >}{< (d_{P-N}(\tau = 0) - \overline{d_{P-N}})^2 >},$$

, where $\overline{d_{P-N}}$ is the average $d_{P-N}$ over all simulation time. The relaxation time $\gamma_{P-N}$ was obtained by fitting $C_{P-N}(\tau)$ to a single-exponential function.

In essence, $C_{P-N}$ illustrates how the conformation of the IDP gradually becomes unrelated to the original one and $\gamma_{P-N}$ characterizes how fast the $C_{P-N}$ decays. To ensure accurate results of $\gamma_{P-N}$ by fitting, we selected the simulation trajectories that are all longer than $10\gamma_{P-N}$.

### References


[1] Roshan Mammen Regy, Wenwei Zheng, and Jeetain Mittal. Using a sequence-specific coarse-grained model for studying protein liquid–liquid phase separation. In *Methods in enzymology*, volume 646, pages 1–17. 2021.

[2] Felipe J Blas, Luis G MacDowell, Enrique de Miguel, and George Jackson. Vapor-liquid interfacial properties of fully flexible lennard-jones chains. *The Journal of chemical physics*, 129(14), 2008.

[3] Aiman Moldagulova and Rosnafisah Bte Sulaiman. Using knn algorithm for classification of textual documents. In *2017 8th international conference on information technology (ICIT)*, pages 665–671. IEEE, 2017.

[4] Xavier Michalet. Mean square displacement analysis of single-particle trajectories with localization error: Brownian motion in an isotropic medium. *Physical Review E*, 82(4):041914, 2010.

[5] Samuel B Alves, Gilson F de Oliveira Jr, Luimar C de Oliveira, Thierry Passerat de Silans, Martine Chevrollier, Marcos Oriá, and Hugo LD de S Cavalcante. Characterization of diffusion processes: Normal and anomalous regimes. *Physica A: Statistical Mechanics and its Applications*, 447:392–401, 2016.

### Additional Figures

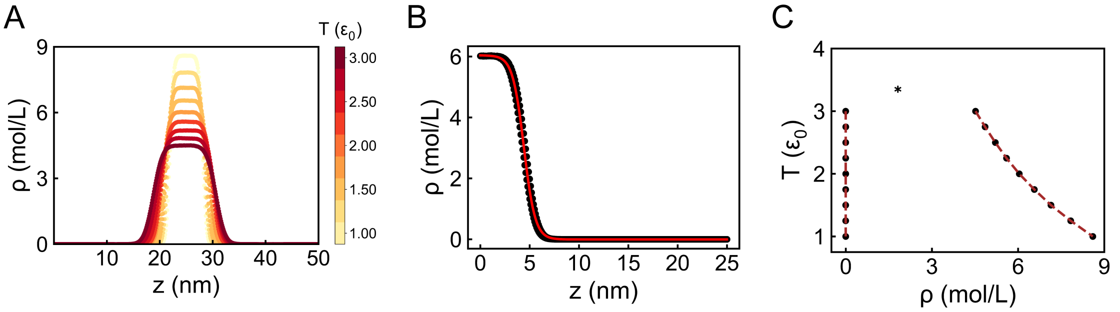


Figure S1: Illustration of fitting of density profile and phase diagram with interaction strength and salt concentration set to be $\varepsilon_{LJ} = 0.20\ \varepsilon_0$ and $C_{salt} = 0.00\ M$, respectively. (A) Density profile along the z-axis varied by temperatures. (B) Fitting of the concentrations ($\rho_z$) at the dense and dilute phases. (C) Phase diagram obtained from the density profile.

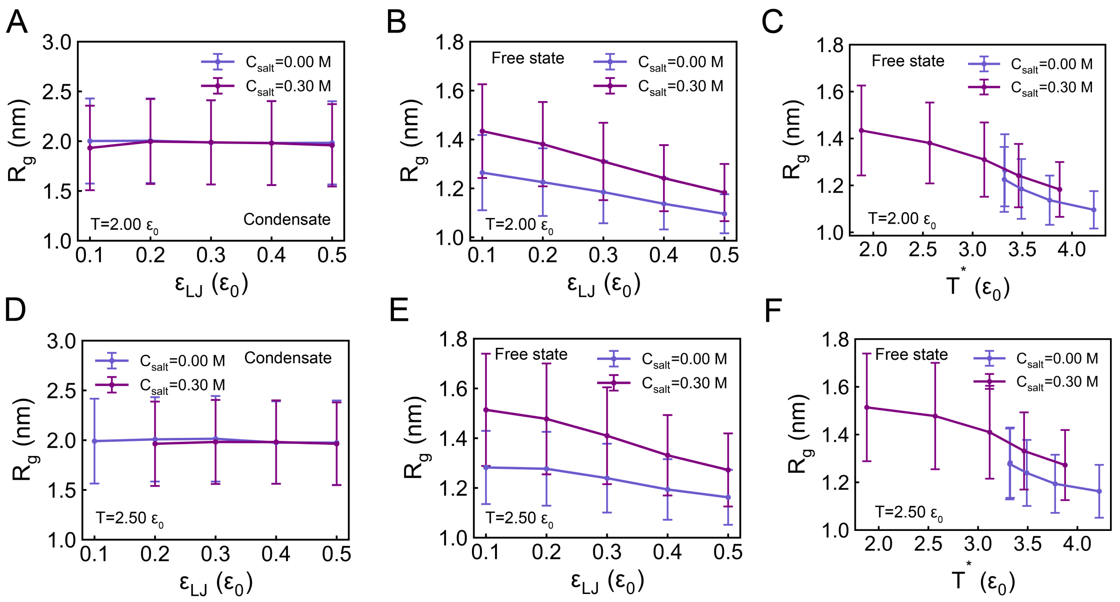


Figure S2: Radius of gyration $R_g$ of the IDP chains at free state and in the condensates at different interaction strengths and environmental conditions. (A) Interaction-dependent behaviors of the average $R_g$ in the condensates at low ($C_{salt} = 0.00\ M$) and high ($C_{salt} = 0.30\ M$) salt concentrations with the temperature set to be $T = 2.00\ \varepsilon_0$. (B) Same as (A) but for the IDP chains at the free state. (C) Average $R_g$ at the free state changes as a function of the critical temperature $T^*$. (D-F) Same as (A-C) but for temperature set to be $T = 2.50\ \varepsilon_0$.

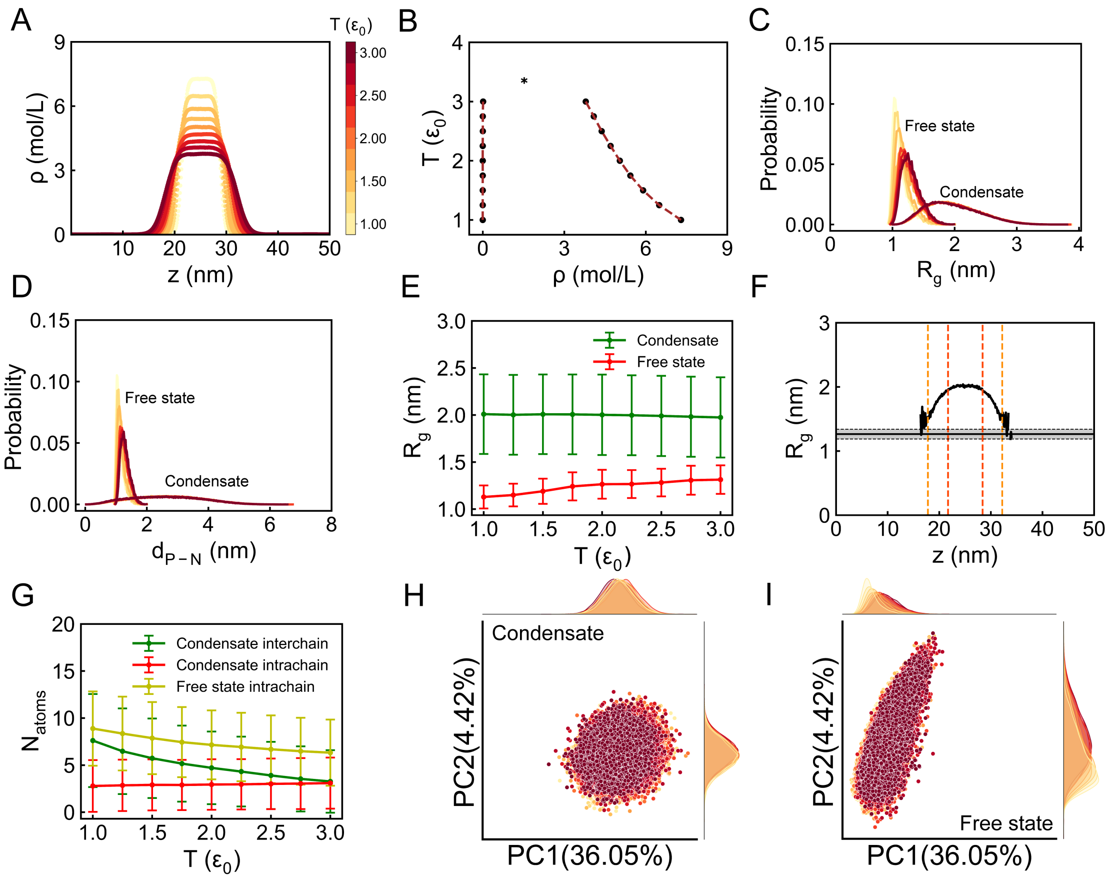


Figure S3: Thermodynamic and conformational properties of IDPs in phase separation. (A) Density profiles along the z-axis varied by temperatures. (B) Phase diagram obtained from the density profile. The critical temperature is denoted as $T^*$. (C) Distribution of the radius of gyration ($R_g$) for IDPs at the free state and in the condensates varied by temperatures, respectively. (D) Distribution of the distance between the positively and negatively charged centers ($d_{P-N}$) for IDPs at the free state and in the condensates varied by temperatures, respectively. (F) Changes of average $R_g$ of IDPs along the z-axis. The red and orange dashed lines represent the boundaries of the center and surface regions of the condensates, respectively. (G) Number of contacts ($N_{atoms}$) for IDPs at the free state and in the condensates. The contacts were further classified into the intra-chain and inter-chain contacts in the condensates. (H, I) Two-dimensional principal component analysis (PCA) of the conformation ensembles of IDPs at the free state and in the condensates, respectively. If not explicitly specified, interaction strength and salt concentration were set to be $\varepsilon_{LJ} = 0.10\ \varepsilon_0$ and $C_{salt} = 0.00\ M$, respectively. Conformational analyses of IDPs were conducted at $T = 2.00\ \varepsilon_0$.

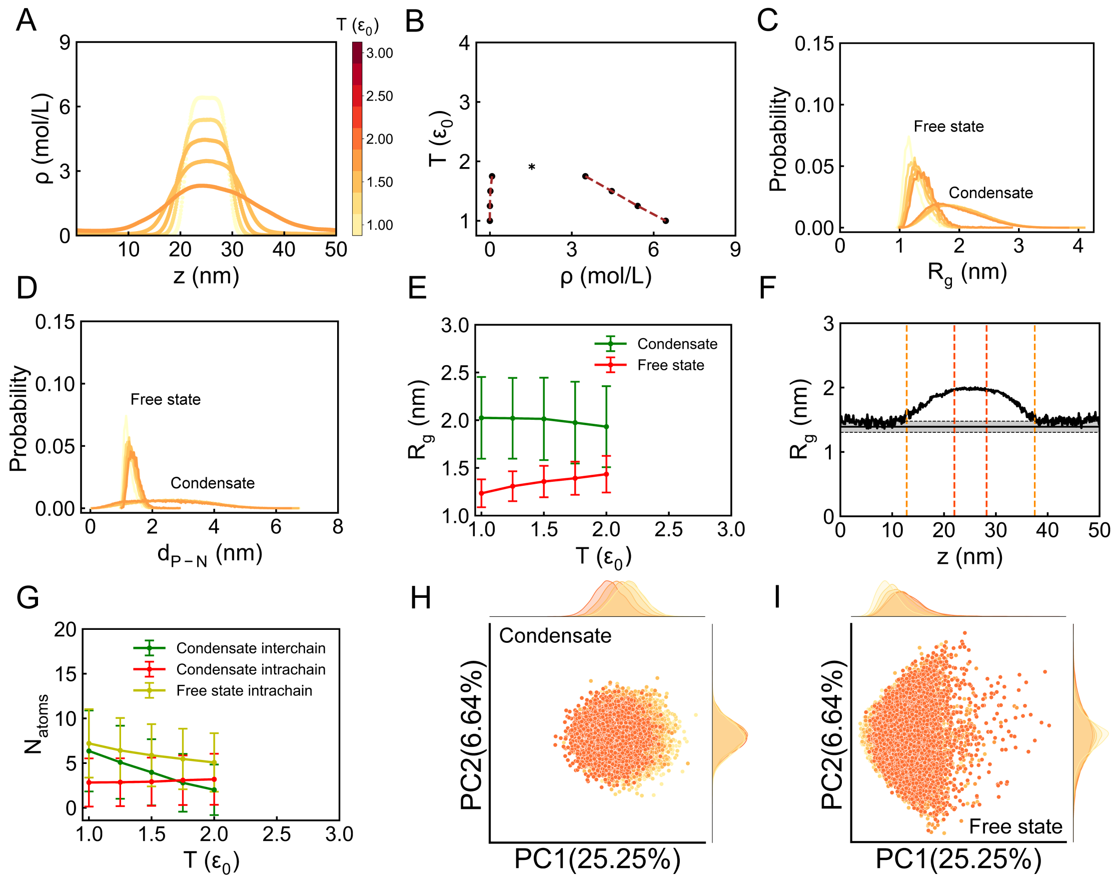


Figure S4: Thermodynamic and conformational properties of IDPs in phase separation. The figure is the same as Figure S3 but for interaction strength and salt concentration set to be $\varepsilon_{LJ} = 0.10\ \varepsilon_0$ and $C_{salt} = 0.30\ M$, respectively.

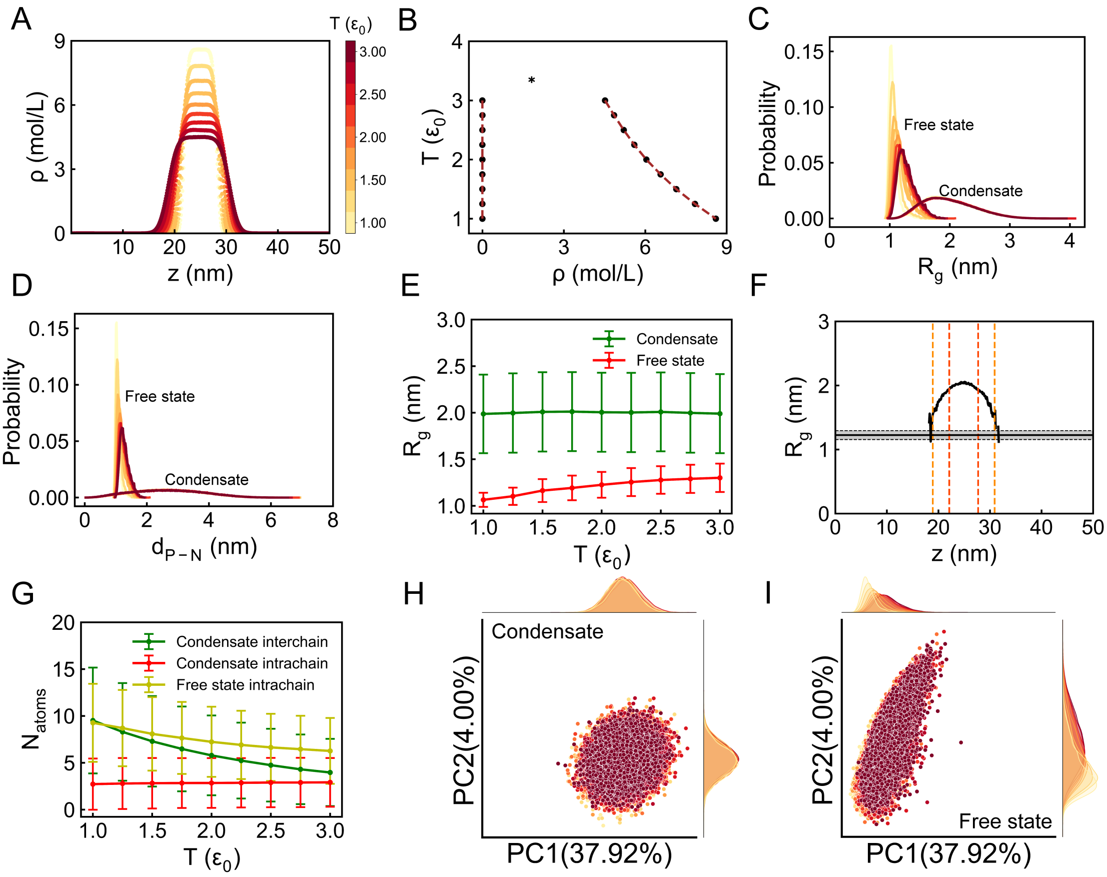


Figure S5: Thermodynamic and conformational properties of IDPs in phase separation. The figure is the same as Figure S3 but for interaction strength and salt concentration set to be $\varepsilon_{LJ} = 0.20\ \varepsilon_0$ and $C_{salt} = 0.00\ M$, respectively. The contents in the figure are also partly shown in Figure 1 in the main text.

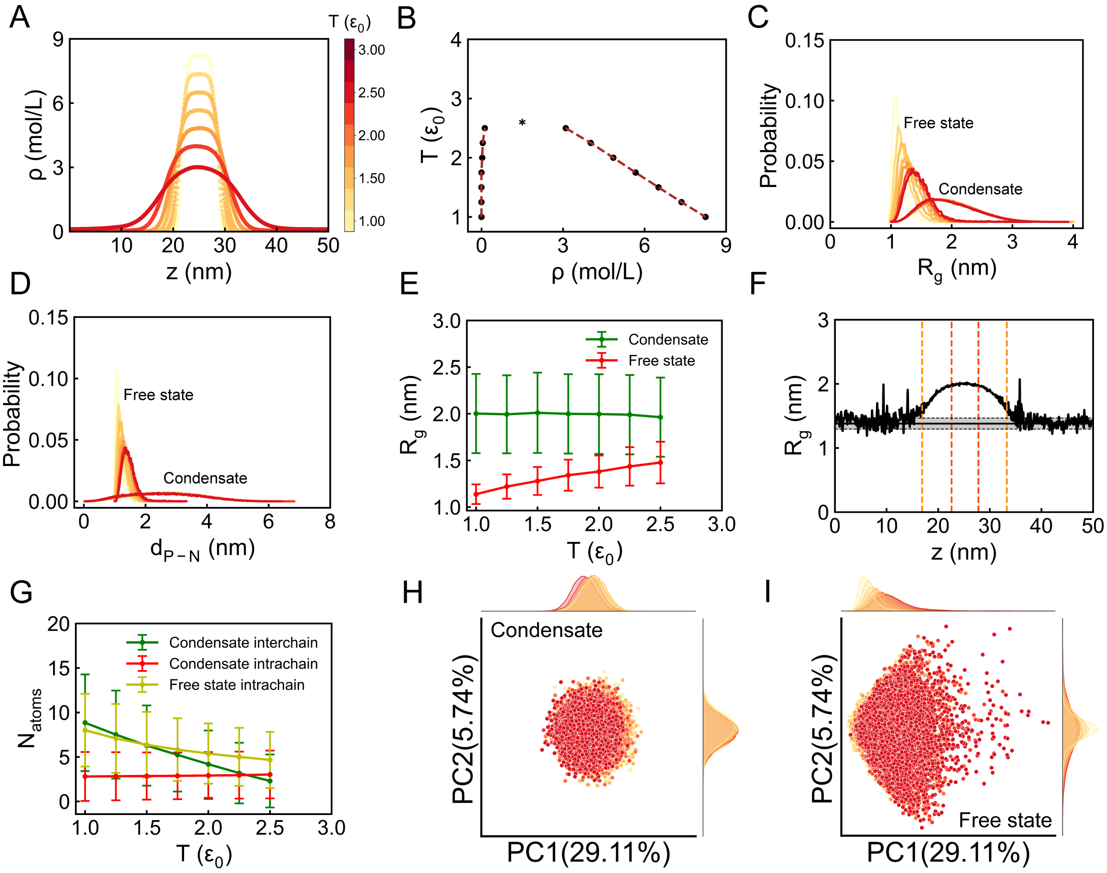


Figure S6: Thermodynamic and conformational properties of IDPs in phase separation. The figure is the same as Figure S3 but for interaction strength and salt concentration set to be $\varepsilon_{LJ} = 0.20\ \varepsilon_0$ and $C_{salt} = 0.30\ M$, respectively.

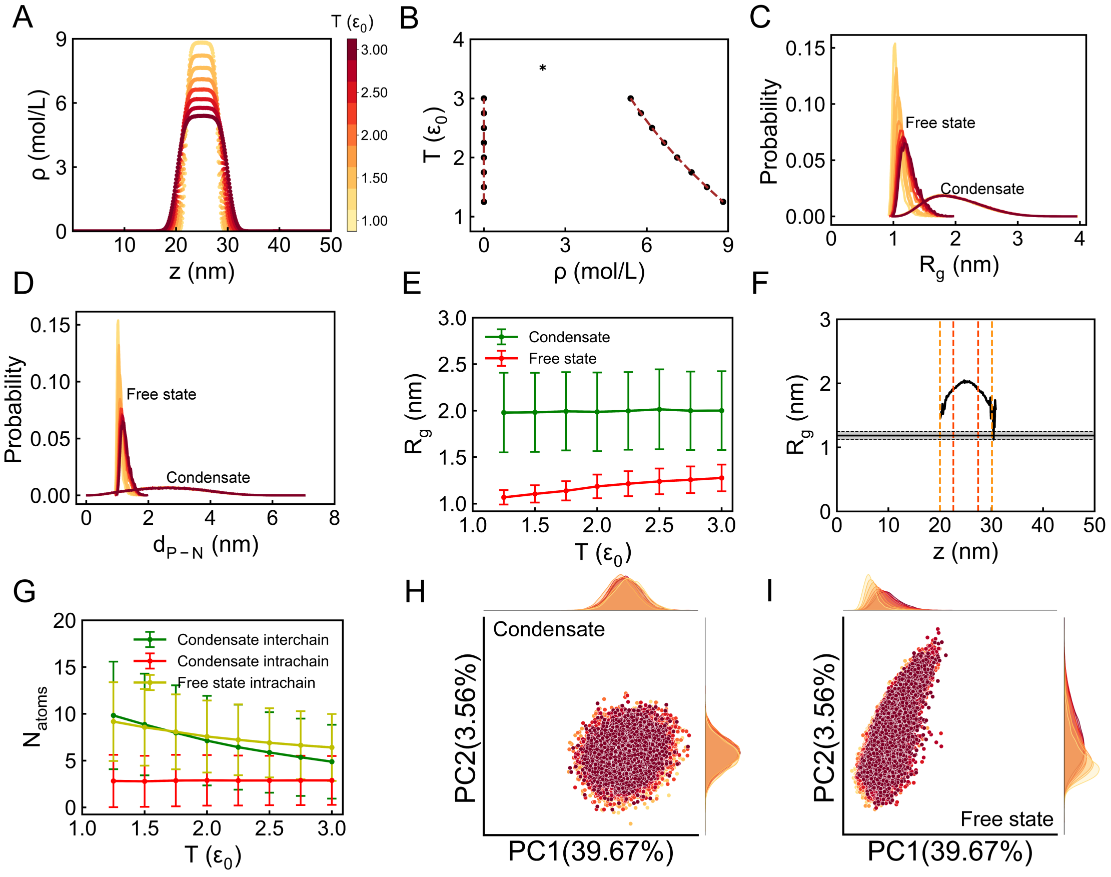


Figure S7: Thermodynamic and conformational properties of IDPs in phase separation. The figure is the same as Figure S3 but for interaction strength and salt concentration set to be $\varepsilon_{LJ} = 0.30\ \varepsilon_0$ and $C_{salt} = 0.00\ M$, respectively.

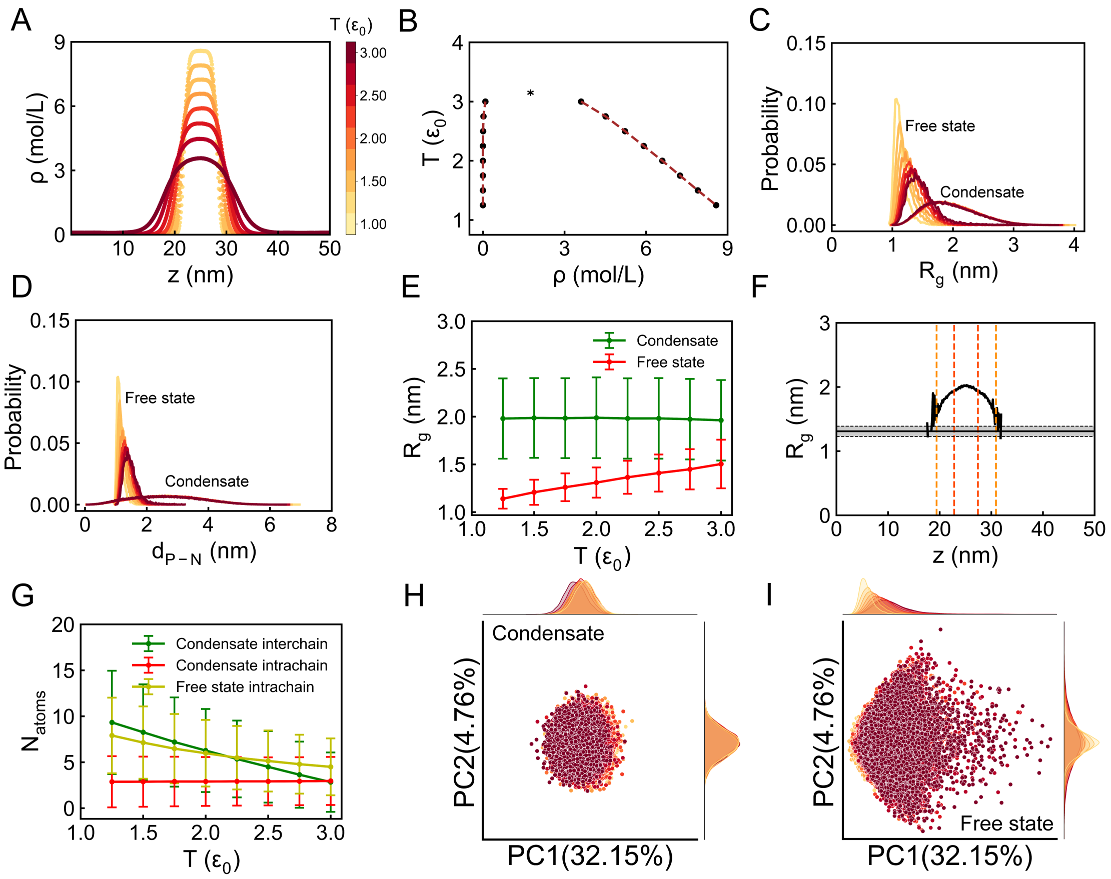


Figure S8: Thermodynamic and conformational properties of IDPs in phase separation. The figure is the same as Figure S3 but for interaction strength and salt concentration set to be $\varepsilon_{LJ} = 0.30\ \varepsilon_0$ and $C_{salt} = 0.30\ M$, respectively.

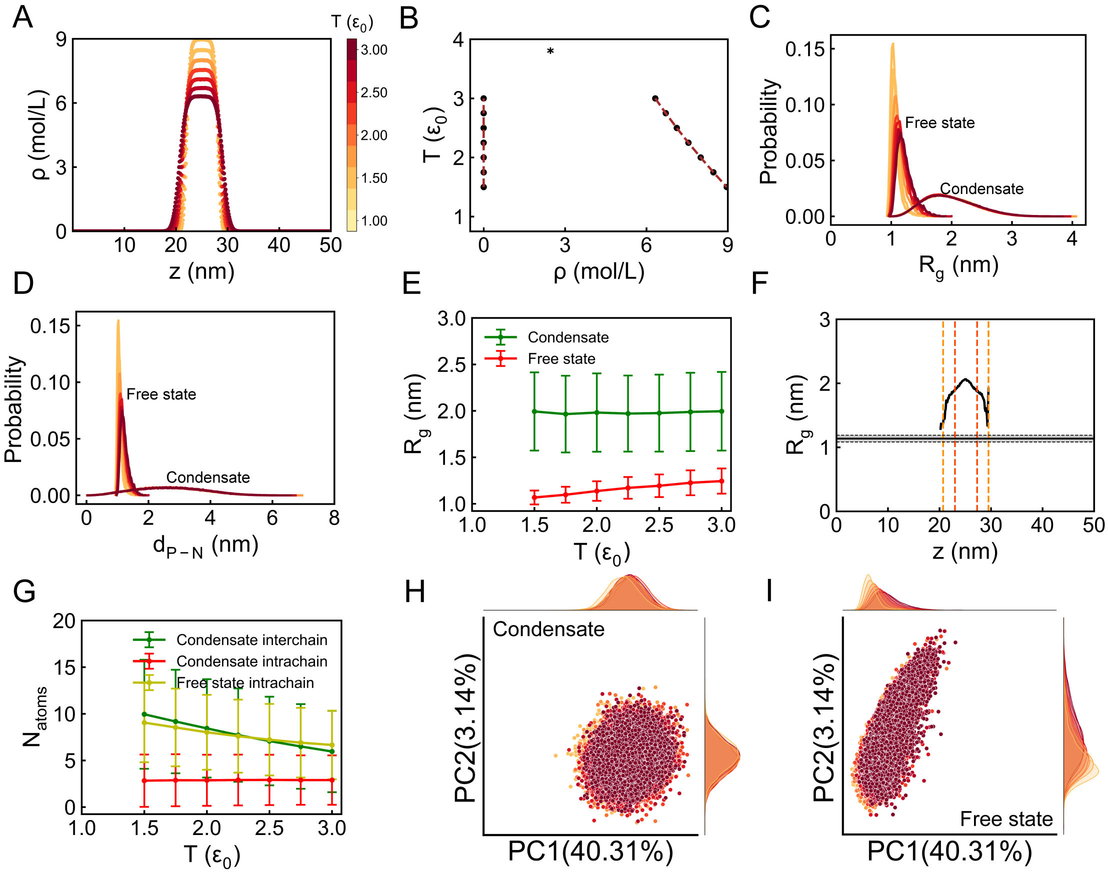


Figure S9: Thermodynamic and conformational properties of IDPs in phase separation. The figure is the same as Figure S3 but for interaction strength and salt concentration set to be $\varepsilon_{LJ} = 0.40\ \varepsilon_0$ and $C_{salt} = 0.00\ M$, respectively.

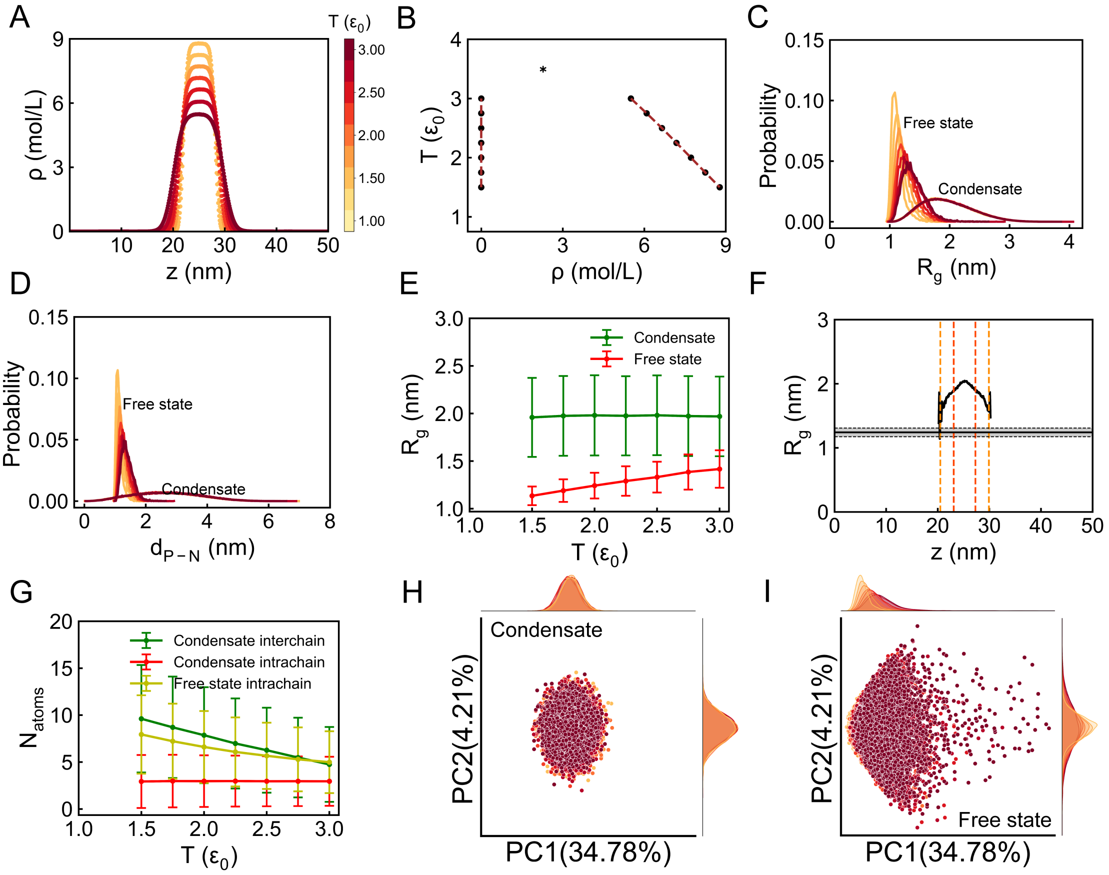


Figure S10: Thermodynamic and conformational properties of IDPs in phase separation. The figure is the same as Figure S3 but for interaction strength and salt concentration set to be $\varepsilon_{LJ} = 0.40\ \varepsilon_0$ and $C_{salt} = 0.30\ M$, respectively.

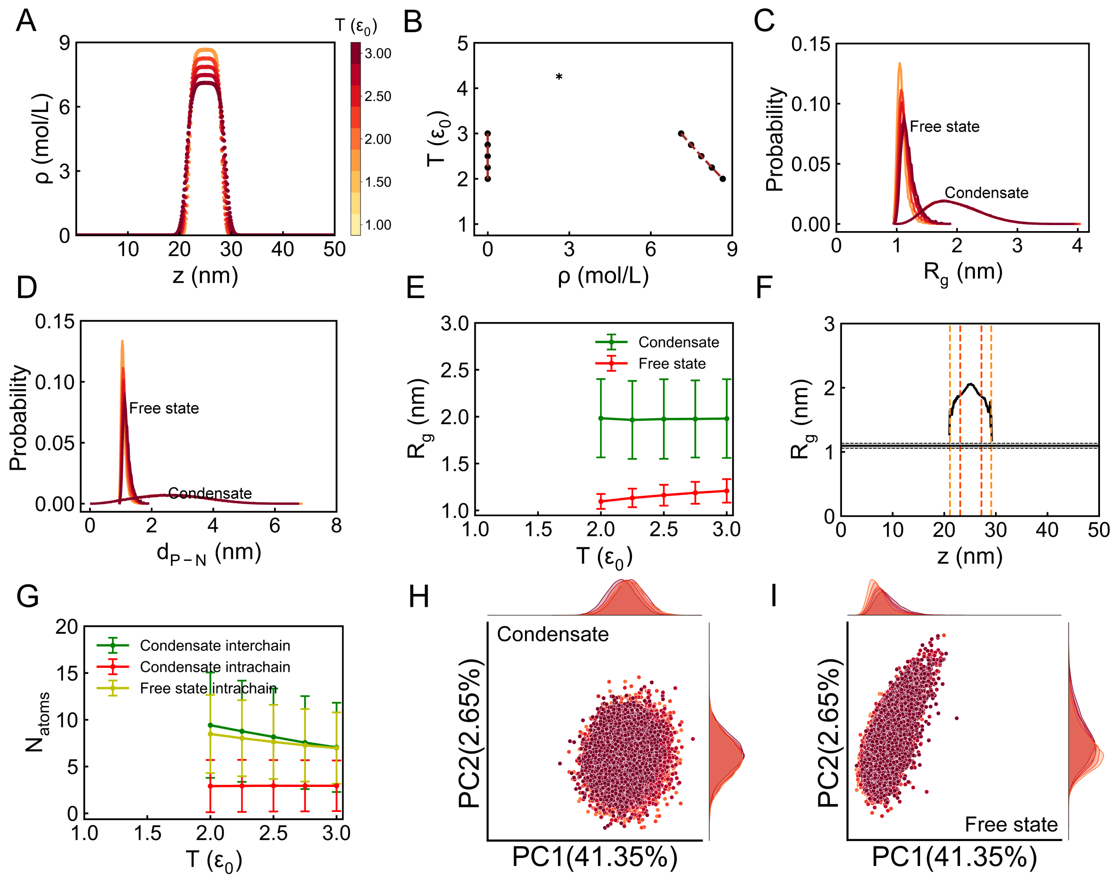


Figure S11: Thermodynamic and conformational properties of IDPs in phase separation. The figure is the same as Figure S3 but for interaction strength and salt concentration set to be $\varepsilon_{LJ} = 0.50\ \varepsilon_0$ and $C_{salt} = 0.00\ M$, respectively.

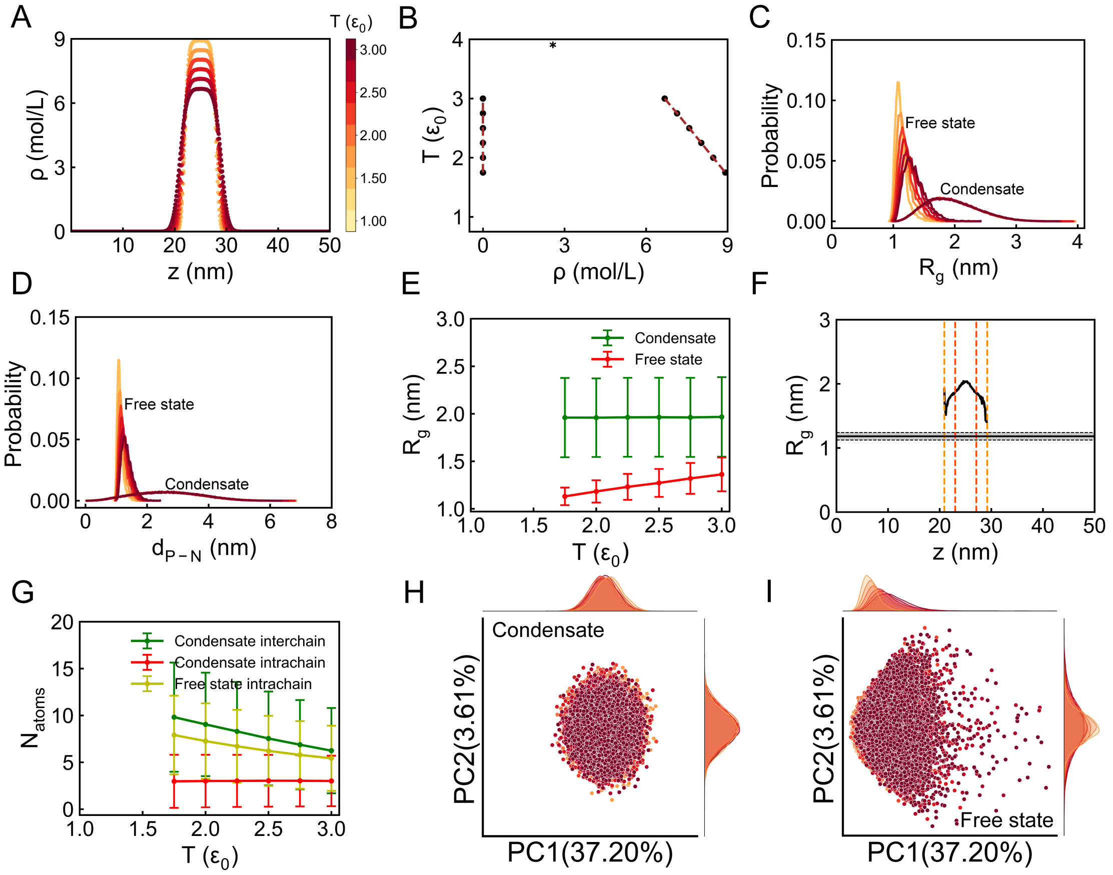


Figure S12: Thermodynamic and conformational properties of IDPs in phase separation. The figure is the same as Figure S3 but for interaction strength and salt concentration set to be $\varepsilon_{LJ} = 0.50\ \varepsilon_0$ and $C_{salt} = 0.30\ M$, respectively.

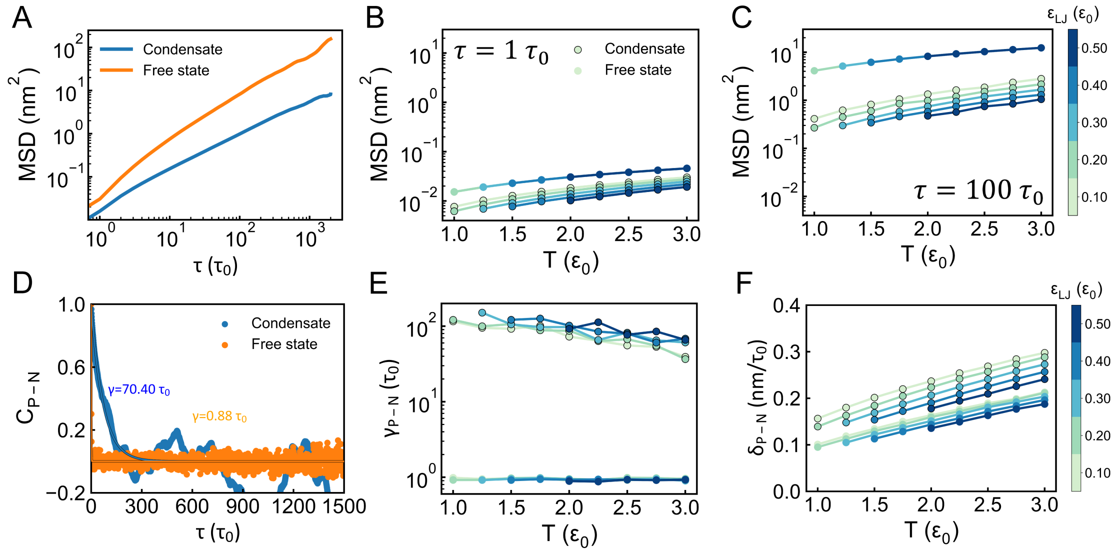


Figure S13: Dynamics of IDPs in the condensates at low salt concentration. The figure is same with Figure 2 in the main text but for salt concentration set to be $C_{salt} = 0.00\ M$. (A) Mean squared displacements (MSD) of IDPs at the free state and in the condensates. (B, C) MSD values at the free state and in the condensates with two time intervals $\tau = 1\ \tau_0$ and 100 $\tau_0$. The dots without the black edges represent the data at the free state and those with edges represent the data in the condensates. The same drawing scheme was applied to the following sub-figures (E) and (F). (D) Autocorrelation of the distance between the positively and negatively charged centers ($d_{P-N}$) in the IDP chain as a function of time. The relaxation time $\gamma_{P-N}$ was obtained by fitting the curve to the single-exponential function. (E) Temperature-dependence of $\gamma_{P-N}$ as a function of $\varepsilon_{LJ}$ at the free state and in the condensates. (F) Rate for describing the IDP conformational dynamics measured by $\delta_{P-N}$, which is the change of $d_{P-N}$ per time unit. The results shown in (A) and (D) were obtained with interaction strength and temperature set to be $\varepsilon_{LJ} = 0.20\ \varepsilon_0$ and $T = 2.00\ \varepsilon_0$, respectively.

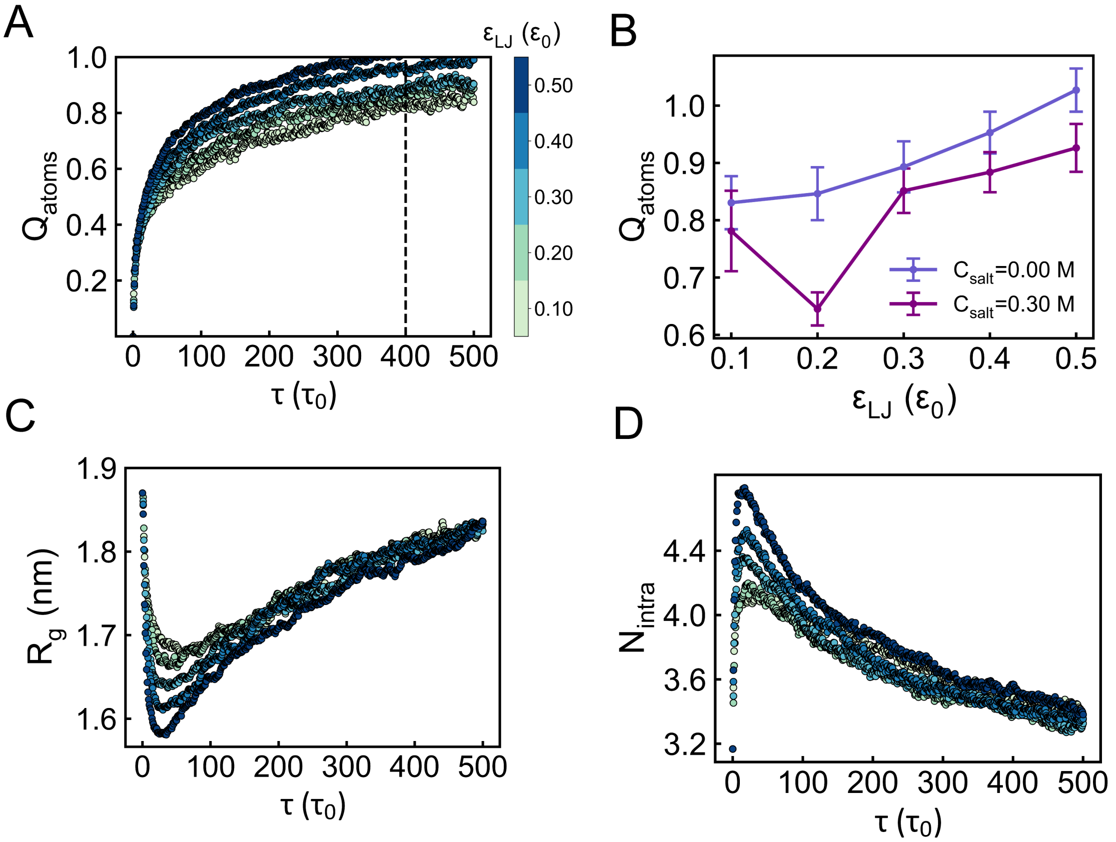


Figure S14: Kinetics of phase separation at low salt concentration. The temperature was set to be $T = 2.00\ \varepsilon_0$, where phase separation can occur steadily at various interaction strengths, as demonstrated in thermodynamic slab simulations. The figure is similar with Figure 3 in the main text but for salt concentration set to be $C_{salt} = 0.00$ $M$. (A) Evolution of phase separation along with time. The process is described by $Q_{atoms}$, which is the fraction of inter-chain contacts formed in the thermodynamic slab simulations. (B) $Q_{atoms}$ values at the time $\tau = 400\ \tau_0$. (C) Evolution of the average $R_g$ in the process of phase separation. (D) Evolution of the average number of intra-chain contacts ($N_{intra}$) in the process of phase separation.
